# Mechanisms of Economic Decisions under Sequential Offers

**DOI:** 10.1101/590422

**Authors:** Sébastien Ballesta, Camillo Padoa-Schioppa

**Affiliations:** Department of Neuroscience, Washington University in St. Louis, St. Louis, MO, 63110, USA; Department of Economics, Washington University in St. Louis, St. Louis, MO, 63110, USA; Department of Biomedical Engineering, Washington University in St. Louis, St. Louis, MO, 63110, USA

**Keywords:** neuroeconomics, orbitofrontal cortex, mutual inhibition, circuit inhibition, monkey, neurophysiology

## Abstract

Binary choices between goods are thought to take place in orbitofrontal cortex (OFC). However, current notions emerged mostly from studies where two offers were presented simultaneously, and other work suggested that choices under sequential offers rely on fundamentally different mechanisms. Here we recorded from the OFC of macaques choosing between two juices offered sequentially. Analyzing neuronal responses across time windows, we discovered different groups of neurons that closely resemble those identified under simultaneous offers, suggesting that decisions in the two modalities are formed in the same neural circuit. Building on this result, we examined four hypotheses on the decision mechanisms. OFC neurons encoded goods and values in a juice-based representation (labeled lines). Contrary to previous assessments, decisions did not involve mutual inhibition between pools of offer value cells. Instead, decisions involved mechanisms of circuit inhibition, whereby each offer value indirectly inhibits neurons encoding the opposite choice outcome. These results reconcile disparate findings and provide a unitary account for the neuronal mechanisms underlying economic decisions.

## Introduction

Recent years have witnessed enormous progress in understanding the neural underpinnings economic choices. This behavior is thought to involve two mental stages – values are assigned to the available options, and a decision is made by comparing values. Evidence from clinical data (Heyman, 2009; Rahman et al., 1999; Volkow and Li, 2004), functional imaging (Bartra et al., 2013; Clithero and Rangel, 2013), neurophysiology (Padoa-Schioppa and Conen, 2017; Rudebeck and Murray, 2014; Schultz, 2015; Wallis, 2012) and lesion studies (Camille et al., 2011; Gallagher et al., 1999; Rudebeck et al., 2013) links economic decisions to the orbitofrontal cortex (OFC). In particular, neurophysiology experiments in which monkeys chose between different juices rewards identified three groups of cells encoding the value of individual options *(offer value),* the binary choice outcome *(chosen juice)* and the *chosen value* (Padoa-Schioppa and Assad, 2006). These variables capture both the input and the output of the decision process, suggesting that these groups of cells constitute the building blocks of a decision circuit. Supporting this proposal, trial-to-trial variability in each cell group correlates with choice variability (Padoa-Schioppa, 2013). Furthermore, neuronal dynamics in OFC reflect an internal deliberation (Rich and Wallis, 2016). Complementing these experimental findings, theoretical work showed that neural networks whose units match the cell groups identified in OFC can generate binary decisions (Fig.1ab) (Friedrich and Lengyel, 2016; Rustichini and Padoa-Schioppa, 2015; Solway and Botvinick, 2012; Song et al., 2017; Zhang et al., 2018). Collectively, these results appear to lay the foundations for a satisfactory understanding of the mechanisms underlying economic decisions. In other words, while many aspects of the circuit depicted in Fig.1b need to be elucidated, the basic scheme would seem to be in place. A fundamental limitation of this assessment is the fact that the vast majority of previous studies examined choices between goods offered simultaneously. Yet, in many real-life decisions, offers appear sequentially. Moreover, in natural settings, subjects often shift their gaze, and thus their mental focus, back and forth between options. Current models for choices under simultaneous offers do not account for choices under sequential offers. More precisely, current models could be modified to do so, but there are multiple ways in which such modification may be done.

**Figure 1.**
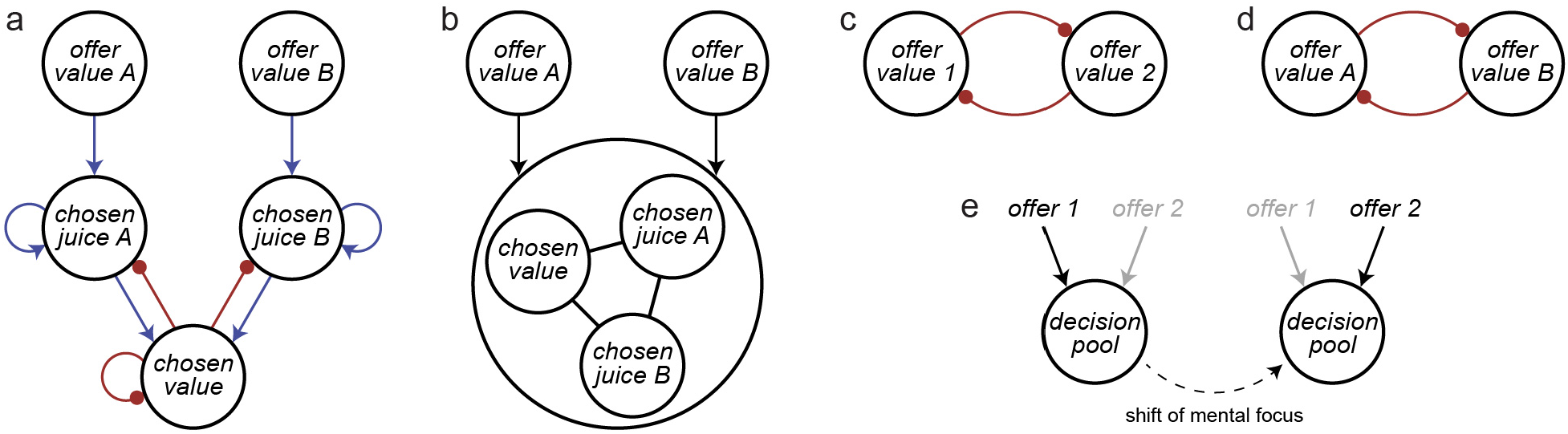
Decision models, **(a)** Biophysically realistic network based on recurrent excitation and pooled inhibition. Blue and red indicate excitation and inhibition, respectively. This model, originally proposed for perceptual decisions (Wang, 2002), captures well the activity of different groups of neurons in OFC during choices under sequential offers (Rustichini and Padoa-Schioppa, 2015). **(b)** General scheme common to multiple models of decisions under simultaneous offers (see main text). In essence, two pools of offer value cells provide the primary input to a neural circuit composed of chosen juice cells and chosen value cells, where decisions are made. The model in panel (a) is one example of the general model shown in panel (b). **(c)** Mutual inhibition at the level of offer values. In this model, decisions take place through mutual inhibition of two populations of neurons encoding the two offer values. This version of the model assumes that options are defined by the sequential order (1 and 2; order-based representation), **(d)** Mutual inhibition in juice-based representation. The model is similar to that in panel (d), except that options are defined by the juice type (A and B). **(e)** Decisions without labeled lines. In this model, the same neural pool examines sequentially the two offers and makes independent accept/reject decisions. Models (c) and (e) are from Hayden and Moreno-Bote (2018).

Importantly, some previous studies did examine choices under sequential offers (Azab and Hayden, 2017, 2018; Blanchard et al., 2015; Blanchard et al., 2017; Strait et al., 2014; Strait et al., 2015; Wallis and Miller, 2003). Unfortunately, their findings are often hard to compare with those summarized above. For one, most studies did not attempt to identify different groups of neurons that might play different roles in the decision process. Furthermore, most analyses assumed that neurons in OFC (or other areas) represent options and values in the reference frame defined by the sequential order, and did not test alternative reference frames (Azab and Hayden, 2017, 2018; Blanchard et al., 2017; Strait et al., 2014; Strait et al., 2015). Failure to consider alternative reference frames can explain negative results obtained when categorization analyses were attempted (Blanchard et al., 2017). In some cases, major conclusions were drawn from negative results (Blanchard et al., 2015). Finally, data sets were relatively small and perhaps insufficient to assess the presence of different cell groups (Azab and Hayden, 2017, 2018; Blanchard et al., 2015; Strait et al., 2014; Strait et al., 2015). Despite these limitations, previous work on choices under sequential offers put forth two testable hypotheses. First, some studies proposed that decisions take place through mechanisms of mutual inhibition between pools of offer value cells (Fig.1cd) (Strait et al., 2014; Strait et al., 2015). Second, other studies challenged the notion that neurons in the decision circuit are associated with individual offers (i.e., the idea of “labeled lines”) (Azab and Hayden, 2017, 2018; Hayden and Moreno-Bote, 2018). Specifically, it was proposed that choices – including binary choices – are processed as sequences of accept/reject decisions (Hayden and Moreno-Bote, 2018). In this view, each offer value is separately compared to an internal benchmark; the decision circuit is constituted of a single pool of neurons that sequentially evaluates and accepts or rejects different options, without a competition between different groups of cells (Fig.1e) (Hayden and Moreno-Bote, 2018). Both proposals are radically in contrast with the scheme depicted in Fig.1b. As a possible reconciliation, some authors speculated that decisions under sequential versus simultaneous offers rely on separate neural circuits (Hayden and Moreno-Bote, 2018; Hunt et al., 2013; Kacelnik et al., 2011; Strait et al., 2015). This last hypothesis, however, has not been tested.

To shed light on the mechanisms of choices under sequential offers, we recorded from a large population of OFC neurons while monkeys chose between two juices offered sequentially and in variable amounts. We analyzed firing rates with approaches similar to those previously used for choices under simultaneous offers (Padoa-Schioppa and Assad, 2006). Most task-related neurons encoded the value or identity of one particular juice type. Their activity in any time window depended on the presentation order. However, an analysis of neuronal responses across time windows revealed the presence of different groups of cells seemingly corresponding to the groups of cells previously identified under simultaneous offers (Padoa-Schioppa, 2013). This result suggested that decisions in the two modalities may be formed in the same neural circuit. Building on this observation, we tested four hypotheses on the decision mechanisms. Our data confuted the notion of a single neuronal pool (Fig.1e). They were also inconsistent with mutual inhibition between pools of offer value cells (Fig.1cd). In contrast, our data pointed to a mechanism of circuit inhibition, whereby neurons encoding the first offer value (input layer) indirectly inhibit neurons representing the second choice outcome (output layer). This inhibited state affects the decision upon presentation of the second offer.

## Results

In the experiments, monkeys chose between two juices offered in variable amounts. The two juices were labeled A and B, with A preferred. The two offers were presented centrally and sequentially (Fig.2a). The terms “offer1” and “offer2” refer to the first and second offer, independently of the juice type and amount. After a delay following offer2, two targets appeared on the two sides of the fixation point, and monkeys indicated their choice with a saccade. For each pair of juice quantities, the sequential order of the two offers varied pseudo-randomly. We were specifically interested in binary choices that require evaluating both offers, with decisions taking place after offer2. In principle, if the value of offer1 is very low or very high, the animal could finalize its decision before offer2. To limit this issue, we designed offer types such that in most trials the animal had to wait for offer2 before making a decision (Fig.2b). In each session, the choice pattern was analyzed with a logistic regression, from which we derived a measure for the relative value of the juices (see Methods).

**Figure 2.**
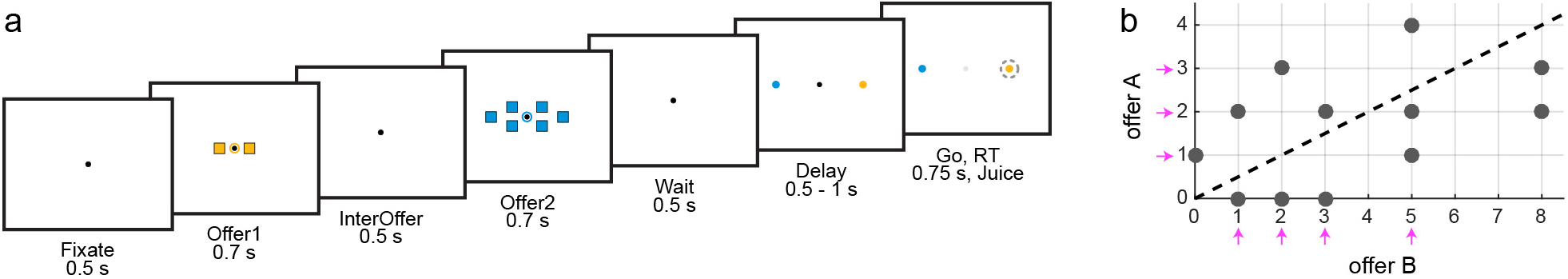
Experimental design, **(a)** The trial started with the animal fixating a center dot. Two offers were presented centrally and sequentially. The color and number of squares represented the juice type and the juice quantity, respectively. In the trial depicted here, the animal chose between two drops of grape juice and six drops of peppermint tea. The second offer was followed by a wait time, after which two saccade targets associated with the two juices appeared on the monitor. After a randomly variable delay, the center fixation point disappeared (go signal), and the animal indicated its choice with a saccade. For each pair of quantities, the sequential order of the offers and the left/right positions of the saccade targets varied pseudo-randomly. **(b)** Offer types for a representative session. The y-axis and x-axis represent quantities of juices A and B offered, gray circles indicate offer types included in the session, and the dotted line (indifference line) captures the relative value of the two juices (ρ = 2). For offers types above (below) the indifference line, the animal usually chose juice A (juice B). Pink arrows indicate juice quantities that, if presented as offerl, prevented the animal from finalizing its decision before offer2 (because, for given offerl, choices were split between the two juices).

### Encoded variables

We recorded the activity of 1267 cells from the OFC of two animals. We examined firing rates in eight 0.5 s time windows aligned with different behavioral events (see Methods). Trials in which juice A was offered first and trials in which juice B was offered first were referred to as “AB trials” and “BA trials”, respectively. An “offer type” was defined by two juice quantities in given order (e.g., [1A:3B] or [3B:1A]); a “trial type” was defined by an offer type and a choice (e.g., [1A:3B, B]); and a “neuronal response” was defined as the activity of one cell in one time window as a function of the trial type. We submitted each neuronal response to an ANOVA (factor: trial type, p<0.001). Neurons passing the criterion in at least one time window were identified as “task-related” and underwent further analysis.

Considering all 8 time windows, 612/1267 (48%) cells were task-related (**Table S1**). Restricting the analysis to the 3 primary time windows (post-offer1, post-offer2, post-juice), 538/1267 (42%) cells were task-related. In a first assessment, many neurons seemed to present different patterns of firing rates in AB versus BA trials. For example, Fig.3ab illustrates the activity of one cell (post-offer1 time window) plotted against variable *offer value 1.* In AB trials, the firing rate increased as a function of the offered value (the cell seemed to encode the *offer value* A). In contrast, in BA trials, the cell was untuned. Such cases were frequent. However, firing rates in the two sets of trials were usually related. For example, for the cell in Fig.3b, the firing rates recorded in BA trials were close to what would be expected if juice A had been offered in quantity 0. In other words, we could define a single variable *offer value A* | *AB* (= *offer value A* in AB trials and = 0 in BA trials) that explained the whole neuronal response. More formally, assuming linear tuning, any neuronal response can be written as:

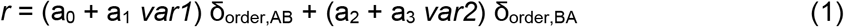

where *r* is the firing rate, *var1* and *var2* are two variables, δ_order,XY_ = 1 if the order is XY and 0 otherwise, and a_0_ … a_3_ are regression coefficients. In the general case, *var1* and *var2* can be any two variables and a_0_ … a_3_ are independent of one another. In contrast, for the vast majority of neuronal responses, we could define a single variable encoded in both sets of trials with the same coefficients, such that Eq.1 reduced to:

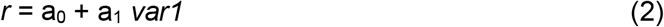

**Figure 3.**
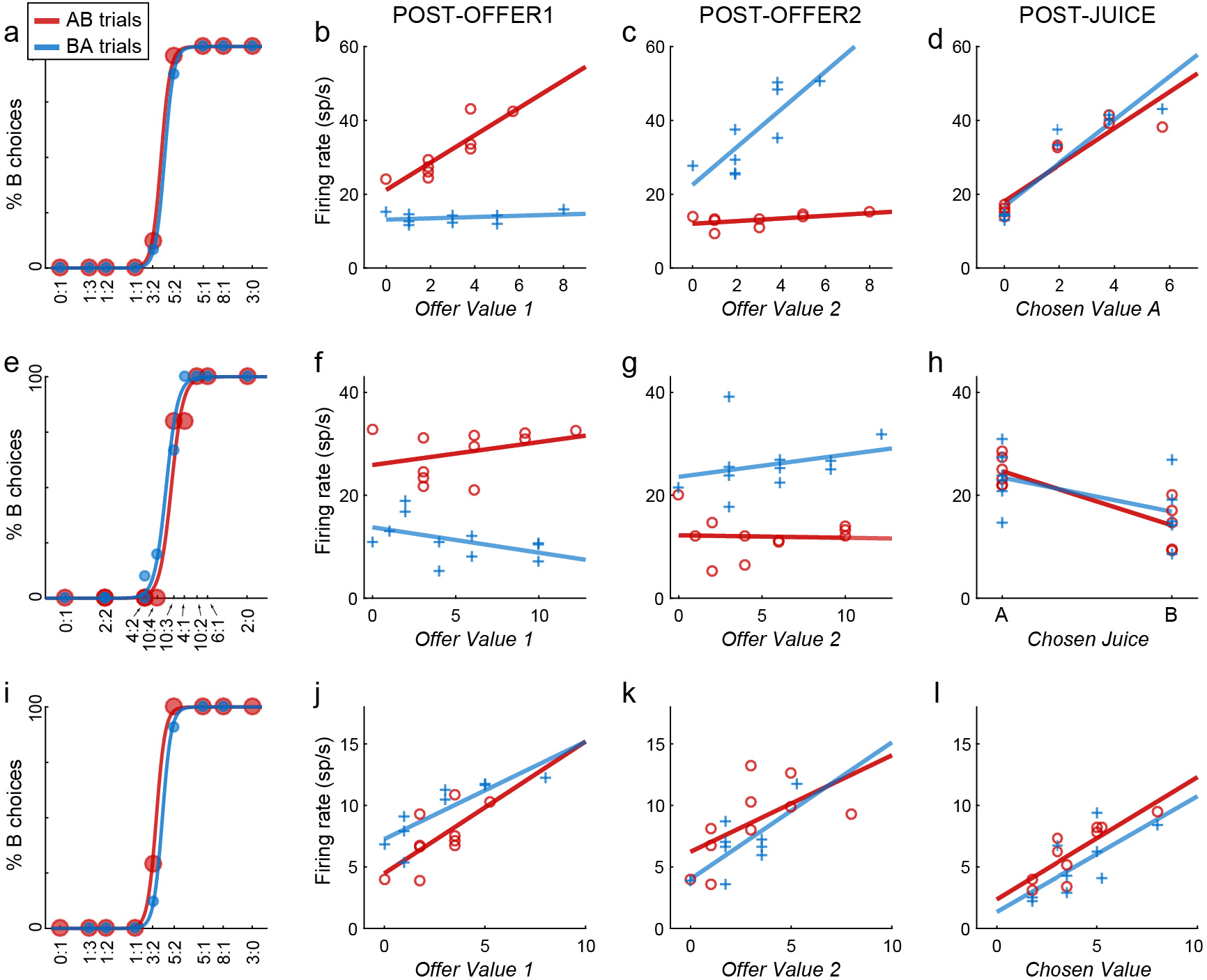
Example cells. Each row illustrates the activity of one neuron. For each cell, the four panels illustrate the behavior and the firing rates in post-offerl, post-offer2 and post-juice time windows. Red and blue colors always refer to AB trials and BA trials, respectively, **(a)** Example cell 1, choice pattern. The percent of B choices is plotted against the log quantity ratio. Sigmoid curves are from a logistic regression (Eq.3). **(b)** Activity in post-offerl Each data point represents one trial type (with firing rate averaged across ~15 trials). Lines were obtained from linear regressions. When juice A was offered (AB trials, red), firing rates increased as a function of the offer value; when juice B was offered (BA trials, blue), firing rates were always low. This neuronal response can be described as encoding the variable offer value A | AB. **(c-d)** In other time windows, this cell encoded variables offer value A | BA (panel c) and chosen value A (panel d). **(e-h)** Example cell 2. In each time windows the activity of this cell was roughly binary. In the post-offerl time window, the activity was high in AB trials (red) and low in BA trials (blue) (variable AB|BA; panel f). In the post-offer2 time window, the activity was high in BA trials and low in AB trials (variable −AB|BA; panel g). In the post-juice time window, the activity was high when the animal chose juice A and low when it chose juice B (variable chosen juice A; panel h). **(i-l)** Example cell 3. In the three time windows, this neuron encoded variables offer value 1, offer value 2 and chosen value.

A description of firing rates according to Eq.2 greatly simplified our understanding.

We considered a large number of variables conceivably encoded in OFC. These included a variable indicating the presentation order (*AB* | *BA*), eight variables associated with individual offers (*offer value A, offer value A* | *AB, offer value A* | *BA, offer value 1, offer value 2,* etc.), two variables capturing the binary choice outcome *(chosen juice, chosen order),* three variables associated with the chosen value (*chosen value, chosen value A*, etc.), and three variables representing the value difference (*value diff (A–B)*, etc.). In total, we examined 18 variables (see Methods, **Table S2**). Each response was separately regressed on each variable. If the regression slope differed significantly from zero (p<0.05), we said that the variable “explained” the response, and we noted the slope sign and the R^2^. Of the 1751 responses passing the ANOVA criterion, 1671 (95%) were explained by at least one of the 18 variables. Fig.4 provides a population summary of the variables encoded in different time windows.

**Figure 4.**
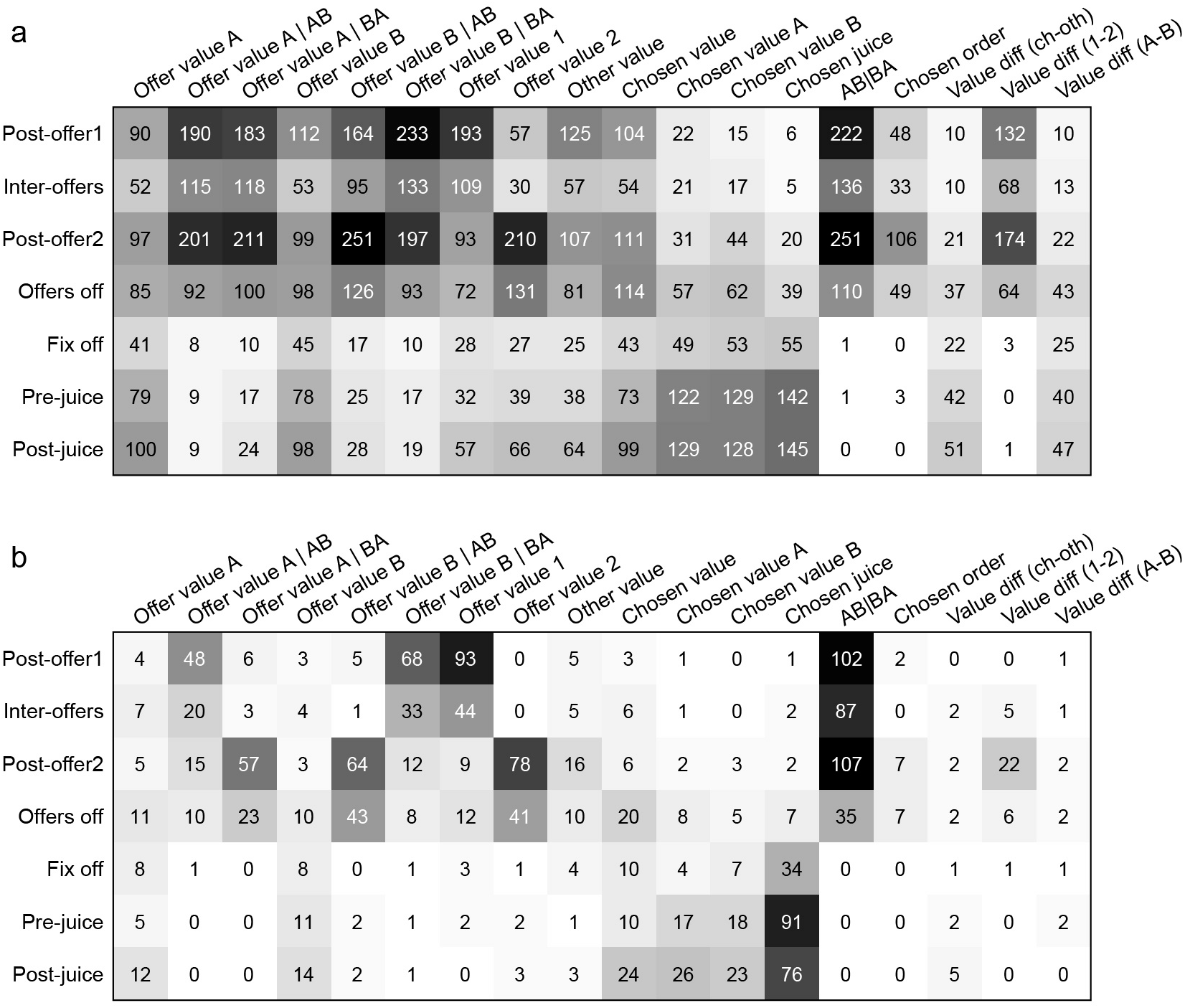
Encoded variables, population analysis, **(a)** Each response passing the ANOVA criterion (p<0.001) was regressed on each of the 18 variables. If the regression slope differed significantly from zero (p<0.05), the variable was said to explain the response. Each response could be explained by >1 variable. Numbers in this panel indicate the number of responses explained by each variable in each time window. For example, variable *offer value A* explained 90 responses in the post-offerl time window. The background image represents the same numbers in gray scale, **(b)** For each response, we identified the variable that provided the best fit (non-zero slope, highest R2). In this panel, numbers indicate responses best explained by each variable in each time window. For example, in the post-offerl time window, variable *offer value A* provided the best explanation for 4 responses. The background image represents the same numbers in gray scale. Notably, different sets of variables explained a disproportionately high number of responses in different time windows. The same cell was often explained by different variables in different time windows, and specific patterns were most frequent. For the analysis of sequences, we focused on time windows post-offerl, post-offer2 and post-juice.

### Neuronal classification

Studies of choices under simultaneous offers found that OFC neurons encoded the same variable across time windows (Padoa-Schioppa, 2013). This property was important because it allowed to identify distinct groups of neurons. In contrast, most neurons recorded here appeared to encode different variables in different time windows. Fig.3a-d illustrates one example. During the post-offer1, the cell encoded the variable *offer value A | AB* (Fig.3b); during the post-offer2, the cell encoded the variable *offer value A | BA* (Fig.3c); during the post-juice, the cell encoded the *chosen value A* (Fig.3d). Another example is shown in Fig.3e-h. In the same three time windows, this neuron encoded variables *AB | BA,* – *AB | BA,* and *chosen juice A.* Finally, in the same time windows, the cell in Fig.3i-l encoded variables *offer value 1, offer value 2,* and *chosen value.* At first, this variability seemed puzzling. However, the variables encoded by a given cell in different time windows were often closely related. For example, consider the neuron in Fig.3a-d. During post-offer1 and post-offer2, the cell encoded the offer value A whenever A was on the monitor. After juice delivery, the cell encoded the value of juice A whenever juice A was delivered. Now consider the cell in Fig.3e-h. The activity of this neuron was roughly binary in every time window. During post-offer1 and post-offer2, the activity was high whenever juice A was on the monitor. During post-juice, the activity was high whenever juice A was delivered. Finally, consider the cell in Fig.3i-l. During post-offer1 and post-offer2, the cell encoded the value of the good present on the monitor. During post-juice, the cell encoded the value chosen (and received) by the animal.

Following these observations, we sought to assess whether OFC neurons typically encode the same *sequence* of variables across time windows. For this analysis, we focused on the 3 primary time windows (post-offer1, post-offer2, post-juice). Considering 18 variables, 2 signs of the encoding and 3 time windows, there were 46,656 possible sequences of variables. Our goal was to assess whether a small subset of sequences could account for the entire data set. In principle, one could conduct an exhaustive analysis considering all the subsets of k = 1,2,3… sequences. For each k, one could identify the best subset as that with the highest explanatory power. Unfortunately, with so many possible sequences, an exhaustive search was not feasible. However, we noticed that sequences providing the best explanation for at least some neuron were relatively few. Thus we focused on 26 sequences that best explained at least 3 cells (**Table S3**), and we conducted an exhaustive search (see Methods). Remarkably, we found that a small number of sequences accounted for most of the population. Specifically, the best subset of 8 sequences explained 510/538 (95%) of task-related cells (Fig.5).

**Figure 5.**
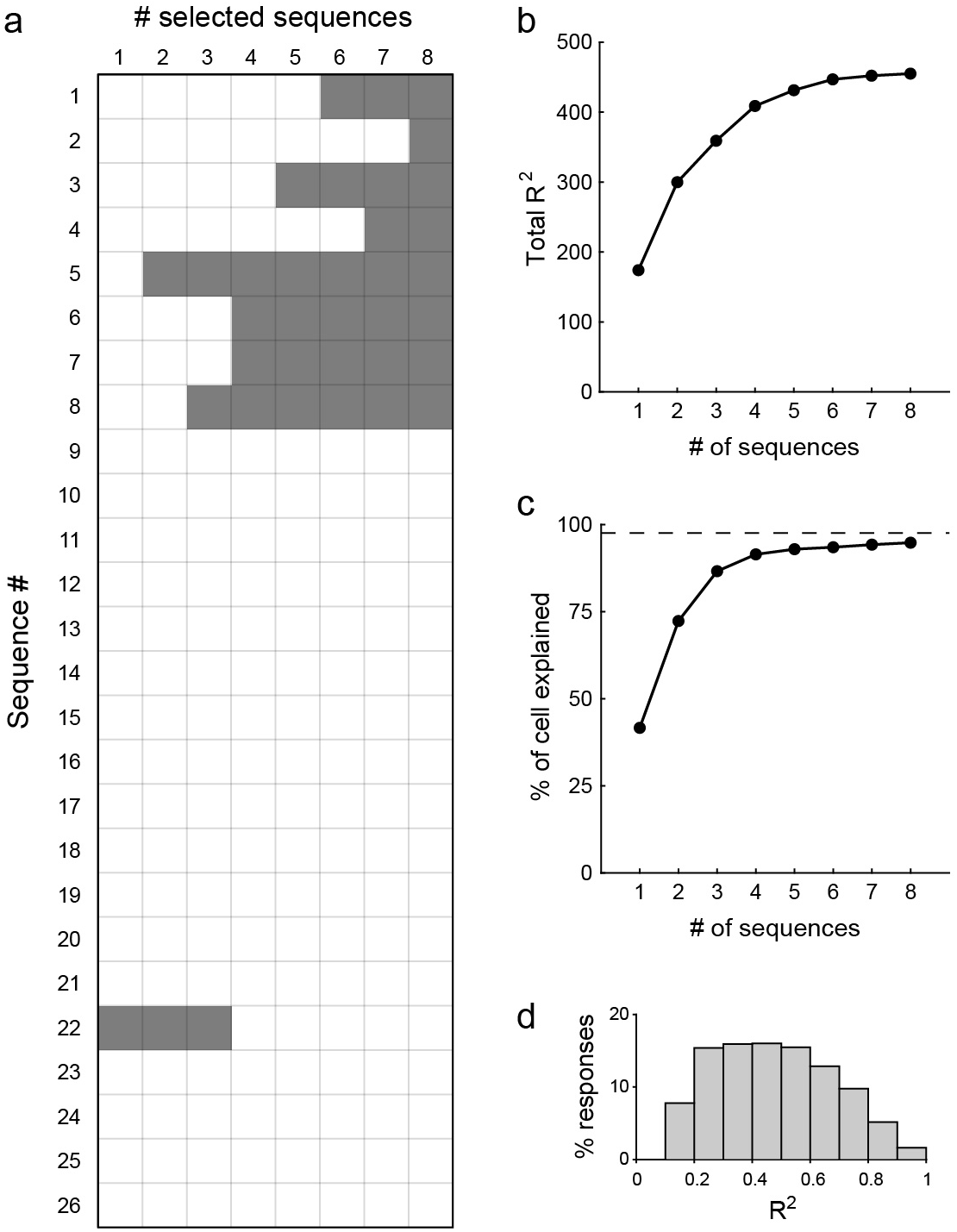
Sequence selection analysis. Given the high number of possible sequences, an exhaustive search for the best subset of k = 1,2,… sequences was prohibitive. Thus we restricted the analysis to sequences that provided the best explanation for ≥3 cells (26 sequences total). Then for k = 1,2,.. we selected the best subset of k sequences based on the total R2 (Methods), **(a)** Sequences selected at each iteration. Sequence numbers 1… 26 (y-axis) correspond to those detailed in Table S3. For each iteration (x-axis), selected sequences are indicated with a gray shade. Notably, in addition to the final 8 sequences, only one other sequence (#22) is ever selected, **(b)** Total R2. The x-axis represents the number of sequences; the y-axis represents the total R2 (summed across time windows and across cells), **(c)** Percent of explained cells. In this panel, 100% corresponds to the number of cells that passed the ANOVA in at least one time window (task-related cells, N=538). The dotted line indicates the percent of cells explained by all 46,656 sequences (N=525, 98%). The best subset of 8 sequences explained 510 cells (i.e., 95% of task-related cells and 97% of cells explained by ail 46,656 sequences). Of note, these 8 sequences explained all of the cells explained by the 26 sequences included in the analysis, **(d)** Distribution of R2 across cells and across variables. For each classified cell (N=510), we considered the time windows that passed the ANOVA, and the R2 obtained from the corresponding variable. The panel illustrates the resulting distribution. Across the population, we measured mean(R2) = 0.48 ± 0.06 (mean ± SEM).

Table 1 details the sequences selected by the best-subset procedure. Notably, these sequences may be divided in 3 groups. Moreover, there seems to be a correspondence between the groups of cells found here and those previously identified during decisions under simultaneous offers (Padoa-Schioppa, 2013; Padoa-Schioppa and Assad, 2006). Specifically, sequences #1, #2, #3 and #4 resemble the cell in Fig.3a-d (considering two juices and two signs of the encoding). These neurons are associated with a single juice and encode the juice value in a linear way. They seem analogous to *offer value* cells identified previously. Sequences #5 and #6 resemble the cell in Fig.3e-h. These neurons are also associated with a single juice, but their activity is binary. They seem analogous to *chosen juice* cells identified previously. Sequences #7 and #8 resemble the cell in Fig.3i-l. These neurons are less easy to interpret.

**Table 1.**
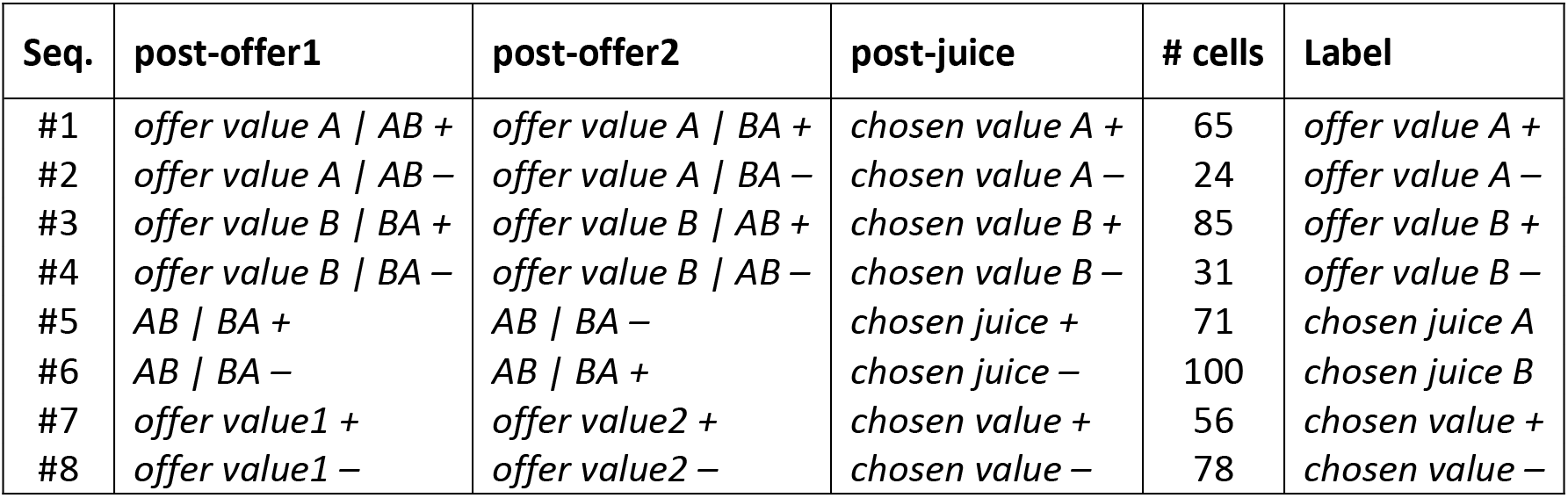
Results of sequence analysis. The sequences shown here are the best subset of 8 sequences. Following the selection procedure, each neuron was assigned to one sequence based on the sequence R^2^ (see Methods). In the table, the leftmost column indicates the sequence number; columns 2-4 indicate the variables encoded in each of the three time windows; column 5 indicates the number of cells assigned to the sequence; the last column indicates the label used to refer to each sequence. In some of the subsequent analyses, we pooled cells from sequences #1 and #3 (*offer value* +), sequences #2 and #4 *(offer value* −), and sequences #5 and #6 (*chosen juice*).

However, there is a possible analogy with *chosen value* cells identified previously, in the sense that these neurons encode the value of either juice, provided that the animal focuses on it. The understanding in terms of mental focus is a valid interpretation of *chosen value* cells in previous studies. Future work will test this correspondence more directly (see Discussion). In the following, we tentatively refer to these groups of cells using the labels *offer value, chosen juice,* and *chosen value*.

### Decision mechanisms

Our aim was to understand how the cell groups detailed in Table 1 support decisions. We specifically examined four hypotheses.

#### Hypothesis 1 (H1): Single pool

It has been proposed that decisions under sequential offers are processed by a single pool of neurons, as sequences of accept/reject decisions, without labeled lines (Fig.1e) (Azab and Hayden, 2017; Hayden and Moreno-Bote, 2018). However, as discussed above, the majority (70%) of task-related cells in our data set were associated with a particular juice, either A or B (sequences #1 to #6). Moreover, these neurons responded differently to offer1 and offer2. These observations are inconsistent with the idea of a single pool and demonstrate the presence of labeled lines.

#### Hypothesis 2 (H2): Mutual inhibition in order-based representation

Several studies examined choices under sequential offers and proposed that decisions take place through mutual inhibition at the level of offer value cells (Fig.1c) (Azab and Hayden, 2017, 2018; Strait et al., 2014; Strait et al., 2015). In those experiments, offers available for choice had some distinctive characteristic represented by a visual trait. For example, in one choice task, the two offers were associated with different reward magnitudes represented by different colors (Strait et al., 2014; Strait et al., 2015). The analysis focused on the post-offer2 time window. Neuronal responses were normalized and regressed against variables *offer value 1* and *offer value 2,* and each term provided a beta coefficient. The key finding was that the two beta coefficients were negatively correlated across the population (beta anticorrelation). This observation was taken as evidence that decisions relied on mutual inhibition at the level of offer values. However, that conclusion was unwarranted because beta anticorrelation may hold regardless of the decision mechanisms. We detail this argument in the **Supplemental Note**. In essence, previous studies assumed that the neuronal representation of goods and values was order-based. However, the choice tasks also afforded a color-based representation. Furthermore, goods defined by the color were offered in different value ranges. In these conditions, neurons encoding the identity or value of individual goods necessarily present beta anticorrelation, independent of the decision mechanisms (**Fig.S1**). Mindful of this issue, we examined beta coefficients in our data set. In neurons recorded with unequal value ranges, we found beta anticorrelation and thus replicated the results of previous studies. However, once differences in value range were controlled for, beta coefficients did not present any positive or negative correlation (Fig.6, **Fig.S2**). Thus we did not find any support for H2.

**Figure 6.**
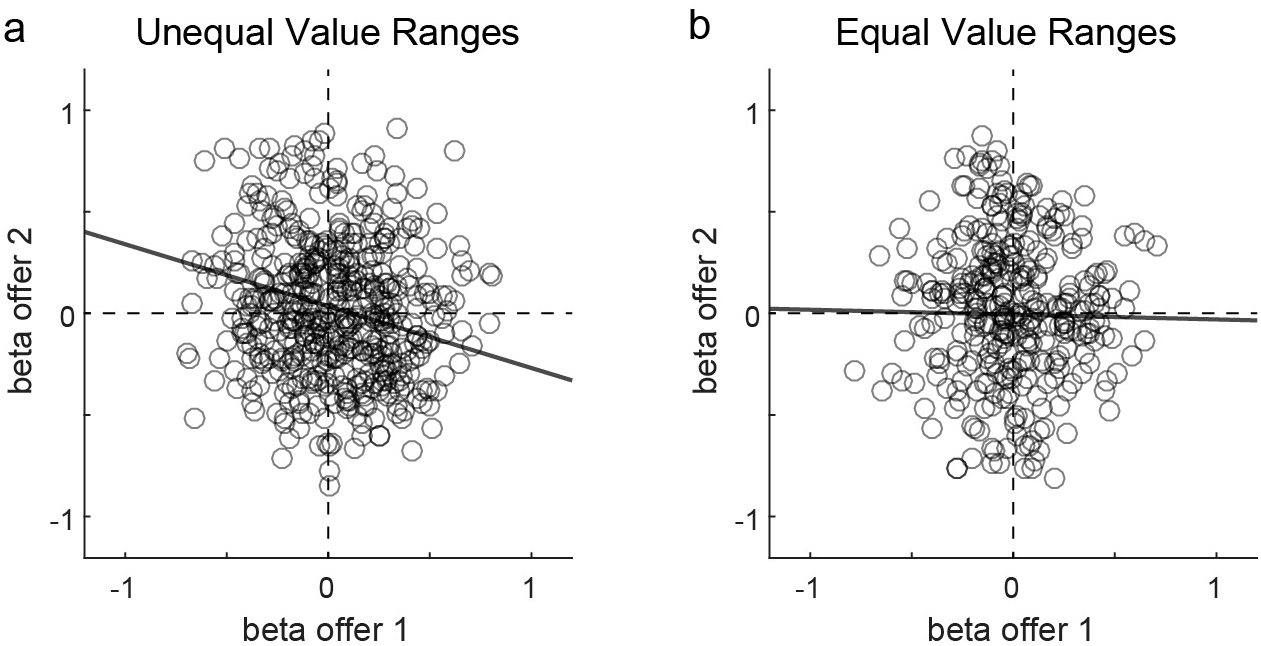
Beta anticorrelation is explained by differences in value range. For each session, ΔV_A_ and ΔV_B_ indicate the value ranges for juices A and B, in common units. In our experiments, value ranges differed from session to session depending on the offered quantities and on the relative value of the juices. Here we divided sessions in two groups depending on whether value ranges were unequal or roughly equal (difference −40%). We then submitted the two populations of neurons to the analysis conducted in previous studies (Strait et al., 2014). The analysis focused on the post-offer2 time window. Firing rates were normalized and regressed against offer valuel and offer value2, which provided two beta coefficients, **(a)** Unequal value ranges (N=505 cells). The two axes correspond to the two beta coefficients and each dot represents one cell. For the population recorded with unequal value ranges, beta coefficients were significantly anticorrelated (corr=-0.11, p<0.02). This result replicates previous findings, (b) Equal value ranges (N=338 cells). For the population recorded with equal value ranges, beta coefficients did not present any positive or negative correlation (corr=-0.02, p=0.78). Importantly, the population in panel **(b)** is substantially larger than that examined in previous studies. Hence, the negative result obtained here does not reflect lower statistical power. For details, see **Supplemental Note**.

#### Hypothesis 3 (H3): Mutual inhibition in juice-based representation

In principle, decisions may entail mutual inhibition between pools of offer value cells associated with different juice types (Fig.1d). If so, when the decision takes place, the activity of neurons directly involved in the decision should reflect the difference between the two offer values. Contrary to this prediction, neurons best explained by the variable *value diff (A-B)* in the post-offer2 time window were vanishingly few (<1%; Fig.4b). Further analysis of the neuronal activity profiles confirmed this point. For each *offer value* cell, we labeled the encoded juice as “E” and the other juice as “O”. We thus refer to EO trials and OE trials depending on whether juice E was offered first or second. (This convention made it possible to pool *offer value A* cells and *offer value B* cells.) We divided trials in three groups depending on whether the value of juice E was low, medium or high. According to H3, the responses of *offer value* cells to juice E should be reduced when juice E is presented as offer2 compared to when juice E is presented as offer1 (due to inhibition). In contrast to this prediction, responses recorded after offer1 (EO trials) and after offer2 (OE trials) were nearly identical and statistically indistinguishable (Fig.7a, **Fig.S3a**). A similar analysis conducted on *chosen value* cells provided similar results (Fig.7b, **Fig.S3b**). In conclusion, our data argue against the hypothesis that economic decisions take place through mutual inhibition at the level of offer values.

**Figure 7.**
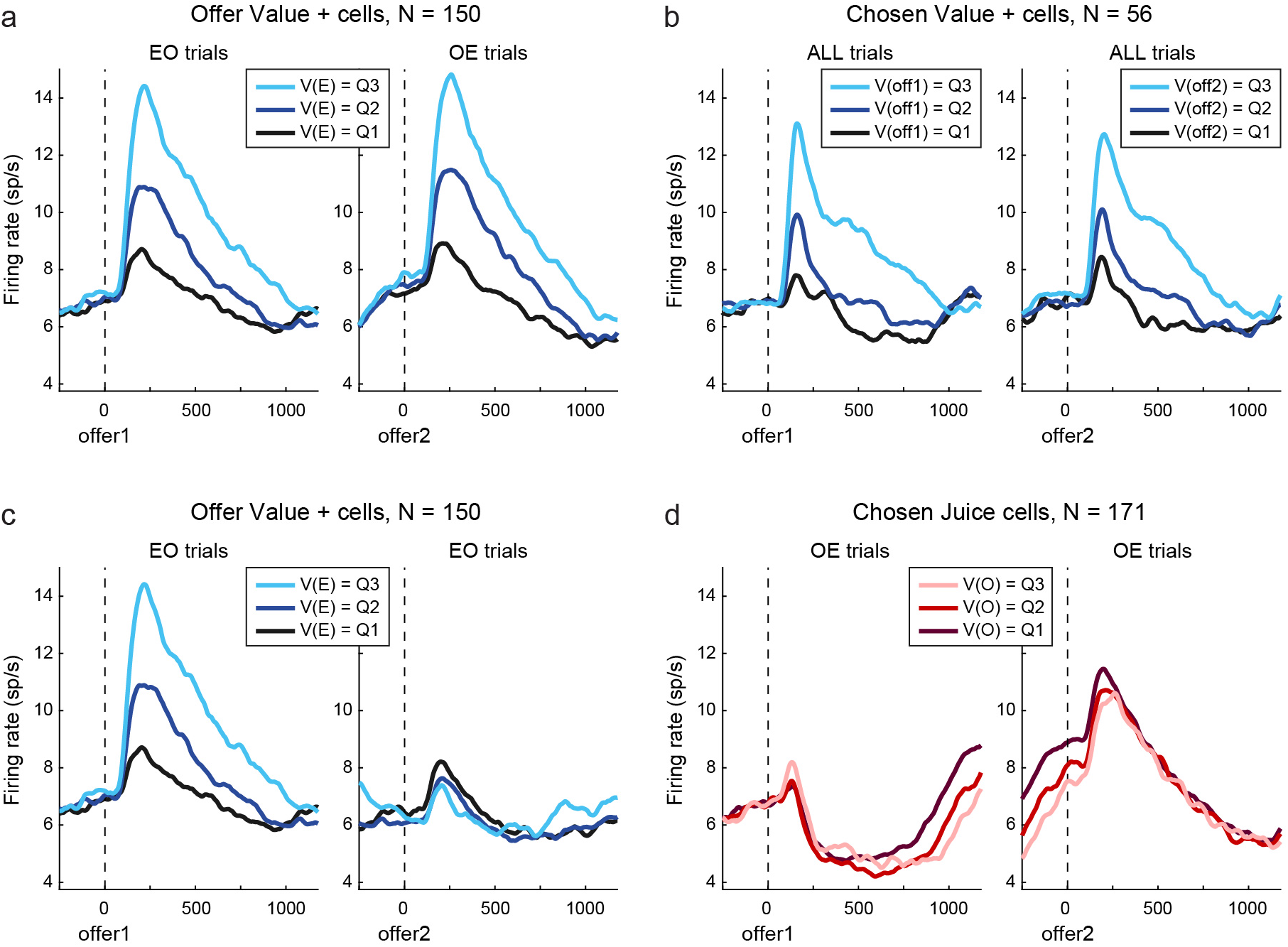
Activity profiles and decision mechanisms, **(a)** Lack of mutual inhibition in *offer value* cells. For each cell, “E” and “O” are the encoded juice and the other juice, respectively. V(E) and V(O) are the value of juice E and the value of juice O, respectively. Left and right panels show the firing rates in EO trials during offerl and the firing rates in OE trials during offer2, respectively. (Hence, both panels illustrate the activity upon presentation of juice E.) For each cell, we divided trials in three groups corresponding to low, medium and high values of V(E) (Q1,Q2,Q3 are tertiles of the distribution for V(E)). We computed the three activity profiles and averaged them across the population. If decisions relied on mutual inhibition between pools of *offer value* cells, activity profiles in the right panel should be substantially lower compared to the left panel (because of inhibition taking place upon offer2, but not upon offerl). In contrast, the two sets of activity profiles were nearly identical, **(b)** Lack of mutual inhibition in *chosen value* cells. Here we pooled trials with different presentation order. In each panel, we divided trials in three groups according to the value offered in the corresponding time window (V(off1) in the left panel, V(off2) in the right panel). Again, the two sets of activity profiles were nearly identical, **(c)** Lack of working memory signal in *offer value* cells. Here we focused on EO trials and we divided trials according to V(E). The left panel is as in panel (a). The right panel shows the same activity traces later in the trial, upon offer2. If decisions relied on a sustained working memory signal encoding the value of offerl until offer2 presentation, traces in the right panel should be separated. The data belie this prediction, **(d)** Negative offset in *chosen juice* cells. Both panels illustrate the activity in OE trials. Trials were divided in three groups according to V(O) (i.e., the value of offerl). Over the course of the delay between the two offers, *chosen juice* cells developed a signal negatively related to V(O). This signal set a negative offset on the activity elicited by offer2 (right panel). The effects illustrated here are quantified in **Fig.S3**.

#### Hypothesis 4: Circuit inhibition

Under H1, H2 or H3, the neural mechanisms underlying decisions under sequential offers would be fundamentally different from those underlying decisions under simultaneous offers. The alternative possibility is that decisions in the two modalities are formed in the same neural circuit. If so, an analysis of choices under sequential offers can provide powerful insights onto the decision circuit.

When offers are sequential, the brain must maintain information about the first offer value and eventually use it in the decision process. Current models for choices under simultaneous offers (Fig.1ab) lack this function and thus do not support choices under sequential offers. However, these models could be adapted to do so. Consider the situation in which offers are sequential and good A is offered first. One reasonable hypothesis is that the pool of *offer value A* cells has a mechanism for working memory. If so, these neurons could maintain a sustained activity throughout the delay intervening between offer1 and offer2. After offer2, the decision would unfold as if the two offers were presented simultaneously. To evaluate this proposal, we focused on *offer value* cells and examined EO trials in both post-offer1 and post-offer2 time windows. Contrary to the prediction, we did not find any sustained activity encoding the value of offer1 (Fig.7c, **Fig.S3c**). An analysis of *chosen value* cells provided similar results (**Fig.S4de**). This observation suggests that the memory trace of offer1 is distributed in the decision circuit, and possibly involves other brain regions.

Aside from the mechanisms of working memory maintenance, we inquired how the value of offer1 might enter the decision process. Our data suggested a mechanism of circuit inhibition whereby *offer value* cells associated with one juice indirectly inhibit *chosen juice* cells associated with the other juice. Fig.7d illustrates the critical finding. In this analysis, we focused on *chosen juice* cells. For each neuron, the juice eliciting higher (lower) firing rates was labeled as juice E (juice O). We examined OE trials, and we divided trials according to the value of the first offer (i.e., the value of juice O). The activity of *chosen juice* cells increased shortly after offer1, and then dropped. However, over the course of the delay intervening between the two offers, the activity gradually increased and became *negatively* modulated by the value of offer1. At the time of offer2, the firing rates of *chosen juice* cells had a strong offset negatively related to the value of the other, non-encoded juice (Fig.7d, **Fig.S3d**).

This observation may be interpreted in relation to the network depicted in Fig.1a. In the model, *offer value* cells and *chosen juice* cells are, respectively, the input and the output layers of the decision circuit. The decision is a competition between attractor states of *chosen juice* cells, mediated by *chosen value* cells. This competition is informed by the input of *offer value* cells and by the initial condition of the network (Rustichini and Padoa-Schioppa, 2015; Wang, 2002; Wong and Wang, 2006). Now consider trials in which juice A is offered first. The activity of *chosen juice B* cells immediately before offer2 represents the network’s initial condition. Our results indicate that *offer value A* cells indirectly inhibit *chosen juice B* cells. By setting a negative offset on the network’s initial condition, this inhibition imposes a bias against juice B (i.e., in favor of juice A) proportional to the offer value A.

The phenomenon illustrated in Fig.7d is reminiscent of the predictive activity previously observed under simultaneous offers (Padoa-Schioppa, 2013) and reproduced by the same network (Rustichini and Padoa-Schioppa, 2015). Under simultaneous offers, the initial condition of the neural assembly, set by the outcome of the previous trial, imposed a bias in favor of repeating the same choice (choice hysteresis). Here, the neural circuit makes a decision only after offer2, and the initial condition is set by offer1. **Fig.S5** illustrates the activity of *chosen juice* cells over the course of a trial.

## Discussion

### A unitary account for economic decisions

There is a broad consensus that binary choices between goods are formed in the OFC (Cisek, 2012; Padoa-Schioppa and Conen, 2017; Rudebeck and Rich, 2018; Rushworth et al., 2012; Wunderlich et al., 2010), but current notions were almost exclusively derived from studies where two offers were presented simultaneously. Yet, in many real-life decisions, offers appear or are examined sequentially. At first, the distinction between choices under simultaneous and sequential offers might seem immaterial. However, computational models for the former do not automatically account for the latter, and previous studies that focused on choices under sequential offers suggested that fundamentally different mechanisms underlie decisions in the two modalities (Hayden and Moreno-Bote, 2018; Hunt et al., 2013; Kacelnik et al., 2011; Strait et al., 2015). Revisiting this pivotal question, we examined the activity of neurons in OFC using a choice task and data analyses comparable to those previously used for choices under simultaneous offers. Remarkably, an analysis of neuronal responses across time windows revealed the presence of different groups of cells seemingly analogous to the groups of cells previously identified under simultaneous offers (Padoa-Schioppa, 2013; Padoa-Schioppa and Assad, 2006). Additional work is necessary to confirm this correspondence. With this caveat, our findings suggest that decisions in the two modalities are formed in the same neural circuit. Building on this result, we tested several hypotheses on the decision mechanisms. Our data clearly argue against the notion of a single pool of neurons (Fig.1e). Contrary to previous assessments, our data also argue against the idea that decisions are made through mutual inhibition between pools of *offer value* cells (Fig.1cd). Conversely, our data suggest that *offer value* cells associated with the first offer indirectly inhibit *chosen juice* cells associated with the second offer – a phenomenon referred to as circuit inhibition.

The idea that the same neural circuit underlies decisions under both simultaneous and sequential offers stands to reason, as two dedicated circuits might seem wasteful. Furthermore, the line between the two modalities is often blurred, because goods presented simultaneously may be examined in sequence. Still, it is worth reflecting on the advantages of circuit inhibition compared to other decision mechanisms discussed above. First, the concept of non-binary accept/reject decisions may be partly misconstrued. Indeed, an accept/reject decision is always a binary choice between some option and aspects of the status quo – often in the form of opportunities – that would be lost if that option were chosen. Future research should examine the neural mechanisms of accept/reject decisions in this perspective. Second, under mutual inhibition, neurons encoding the two offer values inhibit each other. As a result, each pool of neurons comes to encode the value difference, or perhaps the maximum between the value difference and zero. From a computational perspective, one drawback of this putative mechanism is that decisions through the calculation of value differences generalize poorly to choices between multiple options. Indeed, the number of differences to be computed increases combinatorially with the number of options, and this large number of operations still does not resolve the decision. In contrast, the dynamic system depicted in Fig.1a generalizes naturally to choices between multiple items (Furman and Wang, 2008; Wang, 2012).

Importantly, the model in Fig.1a presents several shortcomings. First, the model does not account for neurons with negative encoding. Second, the model does not incorporate the working memory function necessary bridge the delay between the two offers. More generally, it remains unclear whether the model in Fig.1a can capture the complex neuronal dynamics observed for different groups of cells in OFC. Hence, more work is necessary to develop a unitary neuro-computational model for economic decisions under simultaneous and sequential offers. Our results indicate the essential components of such model.

### Implications for choices under free viewing

Although this was cast as a study of choices under sequential offers, the results also shed light on another fundamental issue, namely the role played by attention or mental focus in economic decisions. It was often noted that neurons in the primate OFC are not spatially selective (Grattan and Glimcher, 2014; Padoa-Schioppa and Assad, 2006). At the same time, recent studies showed that neurons in this area can be modulated by gaze or attentional shifts (McGinty et al., 2016; Xie et al., 2018). Our results corroborate both of these observations, as many neurons encoded the value or identity of a particular juice, but only when that juice was present on the monitor. Of note, our animals maintained center fixation while offers appeared and disappeared centrally on the monitor. Thus our results indicate that neurons in OFC are modulated by shifts of mental focus, not by shifts of gaze direction per se. This understanding is completely consistent with previous findings (Krajbich et al., 2010; McGinty et al., 2016; Xie et al., 2018).

From the perspective of the choice system, situations in which subjects shift their gaze or the attention spotlight back and forth between two options on display are presumably similar to the situation examined here, where the gaze is fixed and two options appear foveally in turn. If so, one might speculate about the neuronal mechanisms underlying decisions in the former case. Our results suggest that, upon each gaze shift, OFC neurons encoding the value of the currently attended offer indirectly inhibit the activity of cells representing the opposite choice outcome. Future studies should test this prediction. If confirmed, this understanding would reconcile seemingly diverging ideas put forth in recent years (Hare et al., 2011; Krajbich et al., 2010; Padoa-Schioppa and Conen, 2017).

### Conclusions

To conclude, we examined the activity of neurons in OFC during choice under sequential offers. An analysis across time windows revealed the presence of different groups of cells seemingly analogous to the cell groups previously identified under simultaneous offers. We thus tested several current hypotheses regarding the decision mechanisms. We found evidence against the proposal that binary choices are processed as sequences of accept/reject decisions. Similarly, our data argued against the idea that decisions rely on mutual inhibition at the level of offer values. In fact, we showed that previous arguments for mutual inhibition were confounded. Conversely, our data suggested that economic decisions entail mechanisms of circuit inhibition whereby cells encoding the value of one offer indirectly inhibit cells encoding the opposite choice outcome. Our results provide a unitary account for decisions under simultaneous and sequential offers and lay the foundations for further modeling.

## Methods

### Experimental design, surgery and recordings

All experimental procedures conformed to the NIH *Guide for the Care and Use of Laboratory Animals* and were approved by the Institutional Animal Care and Use Committee (IACUC) at Washington University. Two male rhesus monkeys participated in the study (G, age 6, 9.6 kg; J, age 7, 10.1 kg). The animals sat in an electrically insulated enclosure with their head restrained. A computer monitor was placed 57 cm in front the animal. The behavioral task was controlled through custom-written software (http://www.monkeylogic.net/). The gaze direction was monitored by an infrared video camera (Eyelink; SR Research) at 1 kHz, with an estimated spatial resolution of 0.2°.

In each session, the animal chose between two juices labeled A and B, with A preferred. The two juices were offered sequentially and in variable amounts. Fig.2 illustrates the task design. Each trial began with the animal fixating a dot (0.35° of visual angle) in the center of the monitor. After 0.5 s, two offers appeared centrally and sequentially. Each offer was represented by a set of colored squares, where the color indicated the juice type and the number of squares indicated the juice amount. For example, in the trial depicted in Fig.2a, the animal chose between two drops of grape juice and six drops of peppermint tea. Along with the offer, a small colored circle (0.75° of visual angle) appeared around the fixation dot. In the case of null offer (0 drops), the circle indicated to the animal the identity of the corresponding juice. The animal maintained center fixation throughout the initial fixation (0.5 s), offer1 time (0.7 s), inter-offer time (0.5 s), offer2 time (0.7 s), wait time (0.5 s), and delay time (0.5-1 s). At the end of the delay, the fixation point was extinguished. The animal indicated its choice with a saccade and maintained peripheral fixation for 0.6 s before juice delivery. Center fixation was imposed with a tolerance 2.5°. In a subset of sessions (37%), offer2 was presented for 0.5 s. Sessions included 300-800 trials and offered quantities varied from trial to trial pseudo-randomly (Fig.2b). For each pair of juice quantities, the presentation order (AB, BA) and the spatial location of the saccade targets varied pseudo-randomly and were counterbalanced across trials. Across sessions, we used 12 different juices, resulting in a large number of juice pairings. The association between juice type and color remained fixed throughout the experiments. The juice quantum (i.e., the volume of one drop) was set between 70 μl and 100 μl of volume and did not change during a given session. Importantly, both animals were initially naive. Both of them were trained directly with sequential offers, without previous exposure to the standard choice task (simultaneous offers).

In each animal, we implanted a head-restraining device and an oval recording chamber (Crist Instruments) under general anesthesia. The chamber (main axes, 50×30 mm) was centered on stereotaxic coordinates (A30, L0), with the longer axis parallel to a coronal plane. Recordings were obtained from individual neurons in the central orbital gyrus of both hemispheres using tungsten electrodes (100 μm shank diameter; FHC) advanced with a custom-made motorized system driven remotely. Electrodes were typically advanced in pairs (one motor for two electrodes), with the two electrodes placed at 1 mm from each other. Electric signals were amplified (gain 10,000), filtered (high-pass cutoff, 300 Hz; low-pass cutoff, 6 kHz; Lynx 8; Neuralynx) and recorded (Power 1401; Cambridge Electronic Design). Action potentials were detected on-line and waveforms (40 kHz sampling rate) were saved to disk for off-line clustering (Spike 2; Cambridge Electronic Design). Only cells that appeared well isolated and stable throughout the session were included in the analysis. The data set thus included 829 cells from monkey G and 438 cells from monkey J, recorded over the course of 209 sessions.

### Analysis of choice patterns

All the analyses were conducted in Matlab (MathWorks Inc). Labels “AB” and “BA” indicate the presentation offer (in AB trials A is offered first). Choice patterns were analyzed with a logistic regression

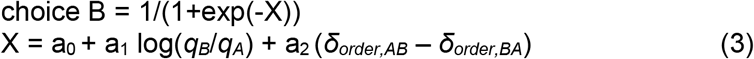

where *q_A_* and *q_B_* are the quantities of juices A and B offered to the animal, δ_*order,AB*_ = 1 in AB trials and 0 in BA trials, and δ_*order,BA*_ = 1 – δ_*order,AB*_· From the fitted parameters, we derived measures for the relative value of the juices (ρ), the sigmoid steepness (η) and the order bias (ε), defined as follows:

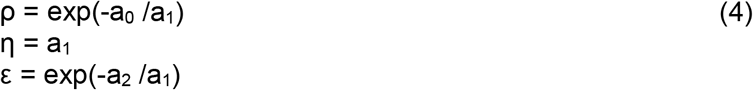

Both animals presented an appreciable order bias favoring offer2 (monkey G: mean(ε) = 0.10, p<0.001; monkey J: mean(ε) = 0.31, p<0.001; t test). However, the order bias was typically small compared to the relative value (monkey G: mean(ρ) = 2.5; monkey J: mean(ρ) = 3.8). Thus the variables examined in the analysis of neuronal data were defined based on ρ, independent of ε (see below).

### Cell classification

Procedures for the analysis of neuronal data were similar to those previously used in studies of choices under simultaneous offers (Padoa-Schioppa, 2013; Padoa-Schioppa and Assad, 2006). Each cell was analyzed in relation to the choice pattern recorded in the same session. We defined eight 0.5 s time windows aligned with different behavioral events: pre-offer (0.5 s before offer1; a control time window), post-offer1 (0.1-0.6 s after offer1), inter-offer (from 0.4 s before to 0.1 s after offer2), post-offer2 (0.1-0.6 s after offer2), offers off (0.6-1.1 s after offer2), fixation off (from 0.1 s before to 0.4 s after extinction of the center fixation point), pre-juice (from 0.4 s before to 0.1 s after juice delivery onset) and post-juice (0.1-0.6 s after juice delivery onset). An “offer type” was defined by two juice quantities in given order (e.g., [1A:3B] or [3B:1A]); a “trial type” was defined by an offer type and a choice (e.g., [1A:3B, B]); and a “neuronal response” was defined as the activity of one cell in one time window as a function of the trial type.

The analysis proceeded in steps. First, data underwent an ANOVA (factor: trial type). Neurons that passed the significance threshold (p<0.001) in at least one time window were identified as “task-related” and included in subsequent analyses. Second, we defined a large number of variables that neurons in the OFC could conceivably encode. These included one binary variable representing the order *(AB | BA*), eight variables representing individual offer values (*offer value A, offer value A* | *AB, offer value A* | *BA, offer value B, offer value B* | *AB, offer value B* | *BA, offer value 1, offer value* 2), variables representing the binary choice outcome *(chosen juice*, *chosen order*), variables capturing variants of the chosen value *(chosen value*, *chosen value A, chosen value B)* and variables capturing the value difference *(value diff (ch–oth); value diff (A–B); value diff (1–2)).* The 18 variables included in the analysis are defined in **Table S2**, and **Fig.S6** illustrates the correlation between variables. Each response passing the ANOVA criterion was regressed on each variable. A variable was said to “explain” the response if the regression slope differed significantly from zero (p<0.05). In this case, we noted the sign of the encoding (regression slope >0 or <0). Each linear regression also provided an R^2^. When a response was explained by more than one variable, the variable with the largest R^2^ was said to provide the best fit for the response. For variables that did not explain the response, we set R^2^=0.

Neurons often encoded different variables in different time windows. However, preliminary observations of neuronal responses across time windows suggested that a small number of variable sequences could account for a large fraction of the population. We thus set up to examine neurons across multiple time windows. Specifically, we focused on the three time windows post-offer1, post-offer2 and post-juice. In this analysis, we used signed variables, where the sign was that obtained from the regression slope. Given 36 signed variables and 3 time windows, there were 46,656 possible sequences. To identify a small number of sequences that can best account for the neuronal population, one would ideally run an exhaustive analysis of all the subsets of k sequences, with k = 1,2,3… However, the large number of possible sequences made this approach computationally unfeasible. To reduce the number of possible sequences, we proceeded as follows. First, for each neuron and for each sequence, we defined the “sequence R^2^” as the sum(R^2^) across time windows. A sequence was said to “explain” a cell if the sequence R^2^ was >0. For each neuron, we identified the sequence that provided the best explanation (highest sequence R^2^). Second, we noted that a relatively small number of sequences (N=387) provided the best explanation for ≥1 cell. Considering sequences providing the best explanation for ≥3 cells along with their mirror sequences (obtained by flipping the encoding signs) further reduced this number to N=26 (see **Table S3**). We thus focused on these sequences, and proceeded with an exhaustive analysis. For k = 1,2,3… we examined each subset of k sequences (>5 × 10^6^ possible subsets). For each subset, we computed the total R^2^ by assigning each neuron to the best sequence in the subset and by summing the sequence R^2^ across all cells. For k = 1,2,3… we thus identified the best subset as that providing the maximum total R^2^. Time windows that did not pass the ANOVA and variables that did not explain a response were normally not considered in these computations. However, these time windows were used to disambiguate cases in which two sequences provided the same sequence R^2^.

Once identified the best subset of 8 sequences, we assigned each neuron to the sequence providing the highest sequence R^2^ (Table 1).

### Analysis of activity profiles

Having identified different groups of neurons, we proceeded with the analysis of their activity profiles. For each *offer value* cell and for each *chosen juice* cell, we labeled the encoded juice as “E” and the other juice as “O”. (For *chosen juice* cells, the encoded juice was that eliciting higher firing rates.) For each cell, we thus refer to EO trials and OE trials depending on whether juice E was offered first or second. These conventions made it possible to pool neurons associated with different juices (A or B). Since we wanted to focus on trials in which the animal could not finalize its decision prior to offer2, we removed from the analysis all forced choices (i.e., trials in which one of the offers was 0). For symmetry, we excluded forced choices independently of whether the null offer was offer1 or offer2. To calculate activity profiles, trials were separately aligned at the times of offer1 and offer2. For each trial, the spike train was smoothed using a kernel that mimicked the post-synaptic potential by exerting influence only forward in time (decay time constant = 20 ms) (So and Stuphorn, 2010). **Fig.S4** was generated with no additional smoothing. For display purposes, we used moving averages of 50 ms in Fig.7 and **Fig.S5**.

For several analyses, we computed the activity profile of a particular population dividing trials in tertiles according to some variable. For example in Fig.7a, we examined *offer value* cells in EO trials and divided trials according to the offer value E (V(E)). To do so, the distribution of V(E) was divided in tertiles for each neuron, and the three activity profiles were averaged across the population. Forced choice were excluded from this analysis.

### Additional references in Supplemental note

(Chau et al., 2014; Hunt et al., 2015; Hunt et al., 2012; Jocham et al., 2012; Louie et al., 2014)

## Acknowledgments

We thank H. Schoknecht for help with animal training and E. Bromberg-Martin, K. Conen, A. Jezzini, I. Monosov, W. Shi and M. Zhang for comments on the manuscript. This research was supported by the National Institutes of Health (grant number R01-DA032758 to CPS).

