## Supplementary material for "Mechanisms of Economic Decisions under Sequential Offers"

### Supplemental Information

**Content:** Supplemental Note, Supplemental Figures S1-S6, Supplemental Tables S1-S3.

### Supplemental note: Mutual inhibition and beta anticorrelation

Numerous studies proposed or were interpreted as showing that decisions take place through mutual inhibition between pools of offer value cells (Hayden and Moreno-Bote, 2018; Hunt and Hayden, 2017). In several cases, the computational models used to fit neural data did not actually involve mutual inhibition at the level of offer values; they involved pooled or indirect inhibition at the level of the choice outcome (as in **Fig.1a**) (Chau et al., 2014; Hunt et al., 2015; Hunt et al., 2012; Jocham et al., 2012; Louie et al., 2014). Other studies, however, examined choices under sequential offers and argued explicitly for mutual inhibition at the level of offer values (**Fig.1c**) (Azab and Hayden, 2017, 2018; Strait et al., 2014; Strait et al., 2015). Their argument was based on a phenomenon referred to as "beta anticorrelation". In the experiments, offers available for choice had some distinctive characteristic represented by a visual trait. For example, in one choice task, two offers were associated with different reward probabilities represented by different colors (Strait et al., 2014; Strait et al., 2015). Neuronal data were recorded from OFC and other brain regions, and the critical analyses focused on the post-offer2 time window. Neuronal responses were normalized and regressed against variables *offer value 1* and *offer value 2*, and each term provided a beta coefficient. The key finding was that the two beta coefficients were negatively correlated across the population. Beta anticorrelation was taken as evidence that decisions relied on mutual inhibition. Here we demonstrate that this conclusion was unwarranted due to a critical confounding factor, namely the difference in value range. We repeated the analysis conducted in previous studies on our data set. Under unequal value ranges, we replicated previous findings (beta anticorrelation). However, once differences in value range were controlled for, beta correlation disappeared. Hence, there was no support for mutual inhibition at the level of offer values.

*<u>Beta anticorrelation does not imply mutual inhibition</u>*

We start with three enlightening examples from our data. The neuronal response in **Fig.S1a-e** was nearly binary – high in AB trials and low in BA trials. In other words, this response encoded the variable *AB|BA* (**Fig.S1b**). Importantly, in this session, the two juices were offered in different value ranges. Expressing both ranges in units of juice B (uB) and indicating with $\Delta V_X$ the value range for juice X, we had $\Delta V_A$=15uB and $\Delta V_B$=6uB. Now consider the same neuronal response plotted against variable *offer value 1* (**Fig.S1c**). Because of the difference in firing rates between AB and BA trials, combined with the difference in value ranges, a linear regression across all data points found a positive slope ($\beta_1>0$). Similarly, a linear regression of all data points against variable *offer value 2* found a negative slope ($\beta_2<0$) (**Fig.S1d**). This response encoded the variable *AB|BA*, but other responses in the same time window encoded the juice identity with the opposite sign (variable –*AB|BA*). For any such response, linear regressions against *offer value 1* and *offer value 2* would find $\beta_1<0$ and $\beta_2>0$. Combining the two groups of responses in a population analysis, these patterns would result in beta anticorrelation.

Notably, the beta anticorrelation emerging from **Fig.S1a-e** is not contingent on this response being recorded in the post-offer2 time window. In fact, if the two value ranges are different, any set of responses encoding variables *AB|BA* and –*AB|BA* in any time window would also show beta anticorrelation. **Fig.S1f-j** illustrates a relevant example. This response, recorded in the post-offer1 time window, encoded the variable –*AB|BA*. Because value ranges were unequal ($\Delta V_A>\Delta V_B$), the regression coefficients had opposite signs (inducing beta anticorrelation in the population). Since the decision could not happen before offer2, an interpretation in terms of mutual inhibition would be paradoxical. Hence, beta anticorrelation does not necessarily imply mutual inhibition.

The two neuronal responses examined so far were binary. However, the argument generalizes to non-binary cases. For example, the response in **Fig.S1kl**, recorded in the post-offer2 time window, encoded the variable *offer value A | BA*. Plotting the response against the variable *offer value 1* confirmed that firing rates did not depend on the value of juice B. However, because $\Delta V_A > \Delta V_B$, a linear regression across all data points found a negative slope ($\beta_1<0$; **Fig.S1m**). Similarly, a linear regression against variable *offer value 2* found a positive slope ($\beta_2>0$; **Fig.S1n**). This neuron was associated with juice A, but other neurons were associated with juice B. Thus in the same time window, other responses encoded the *offer value B | AB*. For any such response, linear regressions against *offer value 1* and *offer value 2* would find $\beta_1>0$ and $\beta_2<0$. Combining all the neurons in a population analysis, these patterns would result in beta anticorrelation.

These examples are directly relevant to previous studies (Azab and Hayden, 2017, 2018; Blanchard et al., 2015; Strait et al., 2014; Strait et al., 2015). In many choice tasks, goods can be represented in multiple reference frame. For example, in our experiments, valid reference frames included that defined by the juice type (A or B, represented by different colors) and that defined by the sequential order (1 or 2). Similarly, in the experiments of Strait et al, valid reference frames included that defined by the reward size (medium or large, represented by different colors[a]) and that defined by the sequential order (1 or 2). The analysis of Strait et al assumed that the representation was order-based. However, this critical assumption was not tested, and it is quite possible that neurons recorded in that study represented goods and values in the reference frame defined by the color[b]. Moreover, the two colors had unequal value ranges. Across trials, the value of the blue option varied uniformly between 0 and 165 μl, while the value of the green option varied uniformly between 0 and 240 μl. This situation is essentially the same as that examined above. In these conditions, neurons that truly encoded the identity of the option presented at any given time (blue or green, in a binary way) would present beta anticorrelation due to the difference in value ranges. Likewise, neurons that truly encoded the offer value of one option (blue or green) would present beta anticorrelation. In either case, beta anticorrelation would hold irrespective of the decision mechanisms and would not imply mutual inhibition.

Other studies that concluded in favor of mutual inhibition presented similar situations (Azab and Hayden, 2017, 2018). One interesting case is the study of Blanchard et al. (2015), where valid reference frames included that defined by the informativeness (low or high, represented by different colors) and that defined by the sequential order (1 or 2). Again, the analysis assumed an order-based representation, but the representation might have been color-based. In this study, the difference in value range was modest (it was essentially equal to the value of information itself). Beta anticorrelation was present but it did not reach significance threshold.

*Population analysis of beta coefficients*

So far, we have shown that differences in value range can induce beta anticorrelation regardless of the decision mechanisms. Next, we examined whether beta anticorrelation holds true once spurious factors are controlled for. To explore this question, we conducted on our data set the same analyses conducted in previous studies. Following Strait et al. (2014), we focused

[a] For simplicity of exposure, we refer only to trials offering these two options (87.5%). However, the presence of trials offering a small sure reward (12.5%) does not affect the argument.

[b] In contrast, our analysis explicitly tested different reference frames. Thus in the present study the statement that different OFC neurons are associated with different juices is a result, not an assumption.

on a 0.5 s time window starting 100 ms before offer2. For each neuron, we normalized firing rates and offer values across all trials. For each trial, we thus obtained normalized measures for the offer values, namely *offer value* $1_{norm}$ and *offer value* $2_{norm}$. We then performed a bilinear regression of the normalized firing rate against *offer value* $1_{norm}$ and *offer value* $2_{norm}$. We thus obtained the two coefficients $\beta_1$ and $\beta_2$, which we plotted against each other for the whole population[c].

We conducted several variants of this analysis. First, we performed the analysis on all the neurons in our data set (all cells, N=1267). Replicating previous findings, beta coefficients were significantly anticorrelated (corr=–0.24, p<0.001; **Fig.S2a**). Second, we noticed that in some of our sessions the offer values for the two juices were correlated with each other (e.g., **Fig.2b**). Given the procedures used in the analysis (normalization and bilinear regression), offer correlation induces a spurious anticorrelation between the beta coefficients. Thus for each session we computed the offer correlation, and we excluded from the data set sessions for which this correlation was statistically significant (p<0.05). Restricting the analysis to the remaining data set (N=843), we found that the beta anticorrelation was lower but statistically significant (corr=–0.07, p=0.036; **Fig.S2b**). For completeness, we repeated the analysis restricting it to sessions presenting significant offer correlation (N=424). As expected, for this population, beta coefficients were strongly anticorrelated (corr=–0.51, p<0.001; **Fig.S2c**)[d]. Third, we noticed that in many of our sessions the two juices were offered in unequal value range. For the reasons discussed above, unequal value ranges induce a spurious anticorrelation between beta coefficients. Thus we restricted the analysis to sessions in which value ranges differed by less than 40% (condition $abs(\log(\Delta V_A/\Delta V_B))<\log(1.4)$). For this data set, (N=338), we did not find any correlation between the beta coefficients (corr=–0.015, p=0.8; **Fig.S2d**) [e]. Conversely, when the analysis was restricted to sessions where value ranges differed by ≥40% (N=505), beta coefficients were significantly anticorrelated (corr=–0.11, p<0.02; **Fig.S2e**). This last result replicates previous findings (Azab and Hayden, 2017, 2018; Strait et al., 2014; Strait et al., 2015).

The key result of these analyses is that shown in **Fig.S2d**. When spurious factors were controlled for, beta coefficients were not (anti)correlated. Note that the population of neurons included in this plot (N=338) is substantially larger than that recorded in any of the previous studies (Azab and Hayden, 2017, 2018; Blanchard et al., 2015; Strait et al., 2014; Strait et al., 2015). Thus the difference between this result and those obtained previously is unlikely due to lack of statistical power in our analysis. More likely, the beta anticorrelation observed in previous studies reflected neurons encoding good identities and/or values in a color-based reference frame, where value ranges were unequal.

In summary, to assess the decision variables encoded in a neuronal population, it is necessary to examine multiple variables defined in multiple reference frames. Failure to do so can lead to erroneous conclusions. Regardless of the decision mechanisms, beta anticorrelation may be induced by various factors such as offer correlation or unequal value ranges. In our data, once

---

[c] The beta coefficients discussed in the previous section ($\beta_1$ and $\beta_2$) were obtained from two linear regressions on non-normalized data. In contrast, the beta coefficients discussed here ($\beta_1$ and $\beta_2$) were obtained from a bilinear regression on normalized data (matching previous studies). We use italics to mark this difference. The arguments discussed in the previous section hold independently of normalization.

[d] In previous studies there was no offer correlation because the two offers were selected independently from uniform distributions.

[e] Further analyses confirmed that the lack of correlation seen in **Fig.S2d** held true even when the analysis was restricted to task-related cells (N=153; corr=–0.07, p=0.38; not shown).

spurious factors were controlled for, beta coefficients did not present any positive or negative correlation. Hence, the results of previous studies were likely due to differences in value ranges.

<u>*Value difference signals*</u>

The examples shown in **Fig.S1** are relevant to another important and closely related issue. Consider the neuronal response in **Fig.S1a-e**. As already noted, due to unequal value ranges, linear regressions against variables *offer value 1* and *offer value 2* found $\beta_1<0$ and $\beta_2>0$. Unequal value ranges also imply that the response is significantly correlated with the value difference (variable *value diff (1-2)*). Indeed, a linear regression against this variable found a significantly positive slope (**Fig.S1e**). Importantly, the response was much better explained as encoding the binary variable *AB|BA* ($R^2=0.76$) than any other variable including the value difference ($R^2=0.27$). Similarly, for the neuronal response in **Fig.S1f-j**, due to unequal value ranges, a linear regression against the value difference (variable *value diff (1-2)*) found a significantly negative slope. The response in **Fig.S1k-o** presents a similar situation.

These examples demonstrate two points. First, if the data analysis was limited to linear regressions against the variable *value diff (1-2)*, one would erroneously conclude that neurons encode the value difference. In contrast, a richer analysis including a large number of variables shows that neuronal data are better explained in terms of juice-related variables (**Table 1**, **Fig.4**). Second, neural activity correlated with the value difference is not necessarily the signature of a decision process. Highlighting this point, such activity can be found even in the post-offer1 time window (**Fig.S1j**), when a decision process can be excluded.

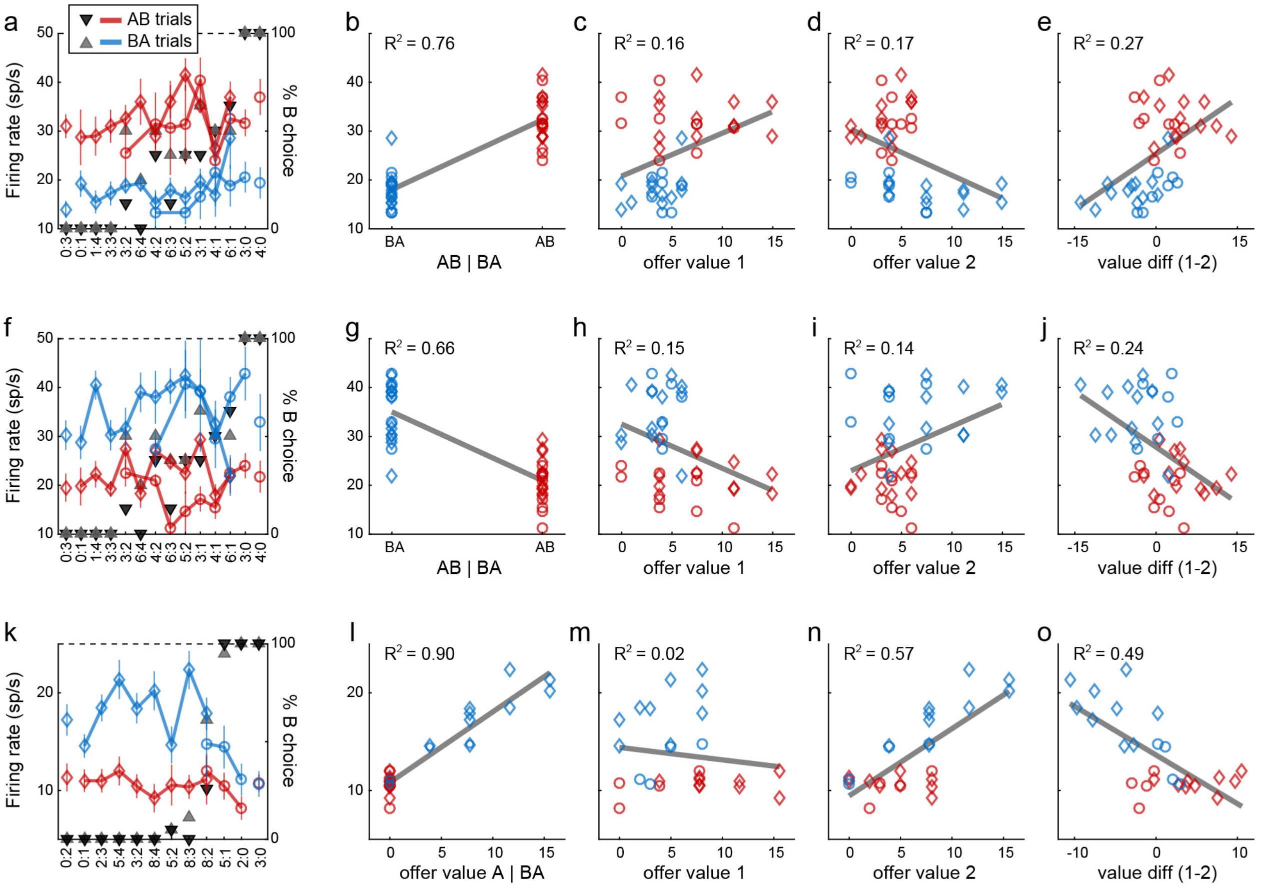


**Figure S1.** Unequal value ranges induce beta anticorrelation. **(a)** Neuronal response encoding the variable *AB|BA*. This panel illustrates the behavior (black and gray triangles) and the neuronal response (color symbols). On the x-axis, offer types are ranked by the quantity ratio $q_B:q_A$. Red and blue colors refer to AB trials and BA trials, respectively. Diamonds and circles refer to trials in which the animal chose juice A and juice B, respectively. This neuronal response was recorded in the post-offer2 time window. Firing rates are roughly binary – high in AB trials (current offer: juice B) and low in BA trials (current offer: juice A). **(b)** Same neuronal response plotted against variable *AB|BA*. Data points are as in panel (a), reordered on the x-axis. The gray line is obtained from a linear regression (R2 = 0.76). **(cde)** Same response plotted against variables *offer value 1*, *offer value 2* and *value diff (1-2)*. In this session, value ranges were unequal, with $\Delta V_A > \Delta V_B$. Thus the range for variable *offer value 1* was larger in AB trials (red) than in BA trials (blue). The neuronal response is still binary. However, a linear regression against variable *offer value 1* (panel (c)) found a positive slope. Similarly, a linear regression against variable *offer value 2* (panel (d)) found a negative slope. Hence, a linear regression against the value difference (panel (e)) found a positive slope. Across a population of cells encoding variables *AB|BA* and *–AB|BA*, this pattern would result in beta anticorrelation. Each panel indicates the R2 obtained from the corresponding linear regression. **(f-j)** Neuronal response encoding variable *–AB|BA*. Same conventions as in panels (a-e). This response was recorded in the post-offer1 time window, before any decision could be made. This example also demonstrates that neural activity correlated with the value difference (panel (j)) does not imply a decision process. **(k-o)** Neuronal response encoding the variable *offer value A | AB*. This response was recorded in the post-offer2 time window. Firing rates varied linearly with the quantity of juice A offered in BA trials (blue); they were low and untuned in AB trials (red). In this session, $\Delta V_A > \Delta V_B$. Although the response was not binary, firing rates were consistently higher in BA trials than in AB trials. Due to the difference in value ranges, linear regressions against *offer value 1* (panel (m)) and *offer value 2* (panel (n)) found negative and positive slopes, respectively.

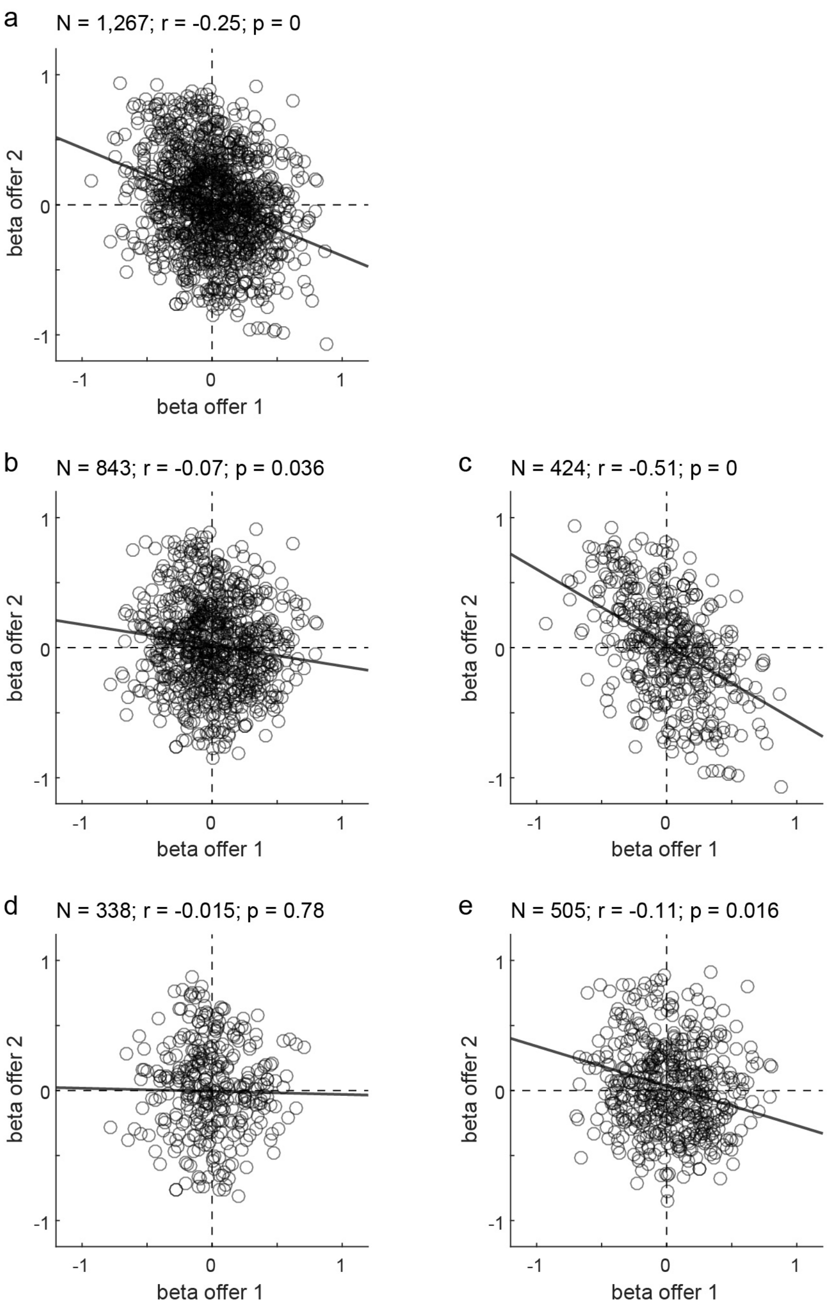


**Figure S2.** Population analysis of beta correlation. **(a)** All cells. **(b)** Cells recorded with uncorrelated offer values. **(c)** Cells recorded with correlated offer values. **(d)** Cells recorded with equal value ranges. **(e)** Cells recorded with unequal value ranges. Each panel indicates the number of cells. In each panel, the r and p values are obtained from Parsons' correlation. The gray solid line is from Deming's regression.

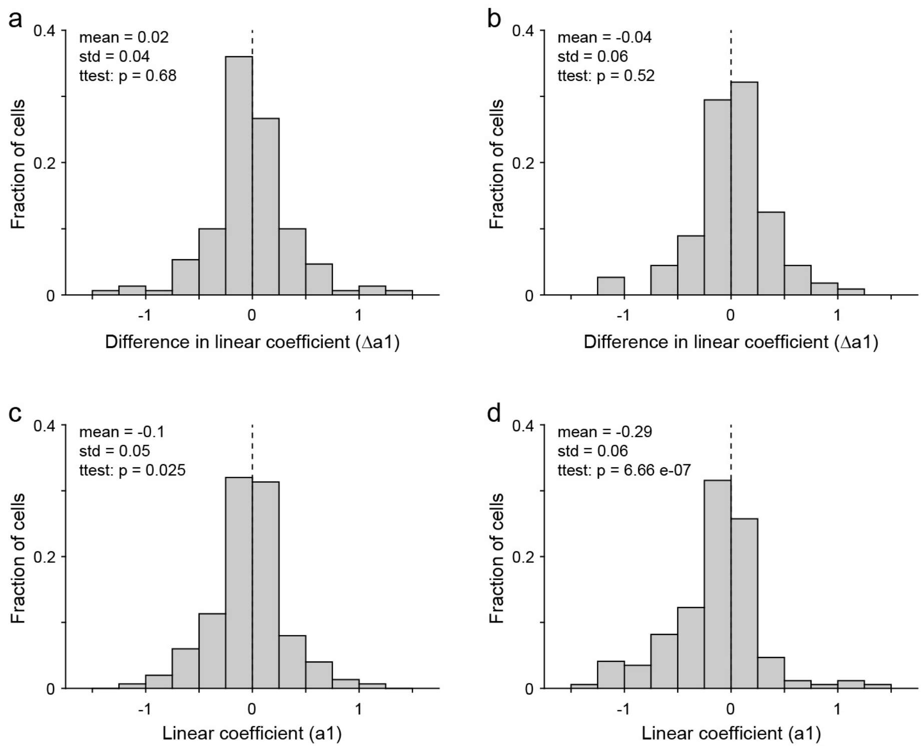


**Figure S3.** Statistical analysis of activity profiles. **(a)** Lack of mutual inhibition in *offer value* cells. The two traces in **Fig.7a** appear nearly identical, suggesting that *offer value* cells respond to the encoded juice (juice E) equally when E is presented first or second. We quantified this point with a regression analysis. For each *offer value* cell, we considered the post-offer1 time window in EO trials and the post-offer2 time window in OE trials. For each group of trials, we performed the linear regression $r = a_0 + a_1 V(E)$, where $r$ is the firing rate. We then compared the two linear coefficients ($a_1$). Across the population, the two measures were statistically indistinguishable (p = 0.68, paired t test). The histogram illustrates the distribution for the difference in $a_1$. **(b)** Lack of mutual inhibition in *chosen value* cells. We conducted a similar analysis on *chosen value* cells. In this case, firing rates in the two time windows were regressed against *offer value 1* and *offer value 2*, respectively. Again, the two linear coefficients were statistically indistinguishable across the population (p = 0.52, paired t test). The histogram illustrates the distribution for the difference in $a_1$. **(c)** Lack of working memory signal in *offer value* cells. The right panel of **Fig.7c** suggests that *offer value* cells encoding the value of offer1 (EO trials) do not represent that signal in a sustained fashion throughout offer2. To quantify this point, we examined EO trials in the post-offer2 time window, and we performed the linear regression $r = a_0 + a_1 V(E)$. We conducted this analysis for each *offer value* cell. The histogram illustrates the distribution of $a_1$. Across the population, linear coefficients were actually negative (mean($a_1$) = –0.1, p = 0.025, t test), although this effect was significant only for one animal. **(d)** Negative offset in *chosen juice* cells. To quantify the effect illustrated in **Fig.7d**, we examined the time window immediately preceding offer2 (from 250 ms before to 50 ms after offer2 onset). For each *chosen juice* cell, we examined OE trials, and we performed the regression $r = a_0 + a_1 V(O)$. The histogram illustrates the distribution of $a_1$. Across the population, linear coefficients were significantly lower than zero (mean($a_1$) = –0.29, p<10-6, t test). This effect held true for both animals (monkey G: mean($a_1$) = –0.30, p = 2 10-5; monkey J: mean($a_1$) = –0.26, p = 0.011; t test).

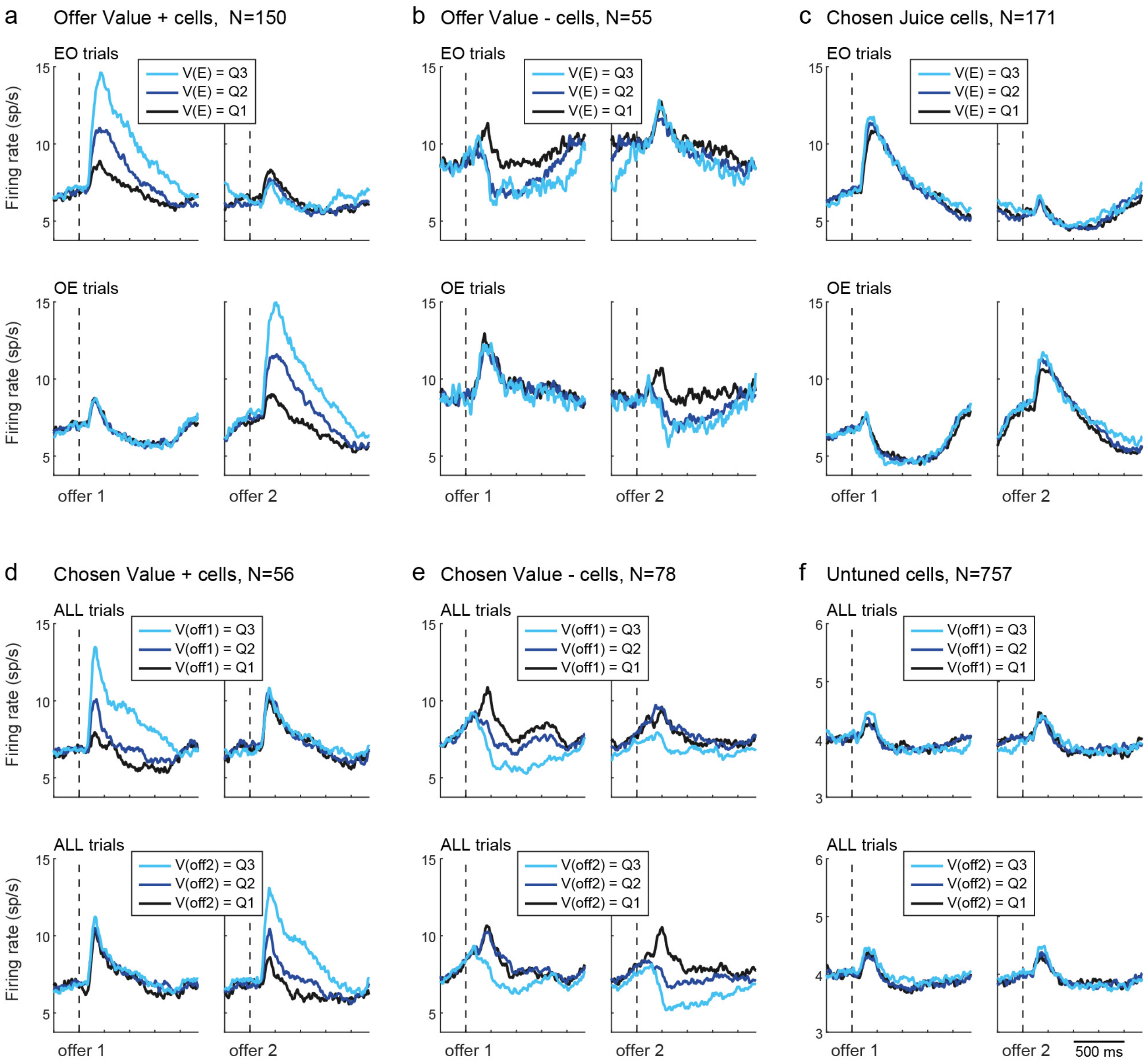


**Figure S4.** Activity profiles, whole population. The figure illustrates the activity profiles for all groups of cells. For each cell group, the activity profiles were aligned separately with offer1 onset and offer2 onset. **(a)** *Offer value* + cells (N=150 cells). E and O refer to the encoded juice and the other juice, respectively. Top and bottom panel illustrate the activity in EO trials and OE trials, respectively. In all panels, trials were sorted according to the value of the encoded juice (V(E)). For each cell, we divided the distribution for V(E) in tertiles (namely Q1, Q2 and Q3) and we computed the activity profiles. We then averaged the activity profiles across the population. **(b)** *Offer value* – cells (N=55). **(c)** *Chosen juice* cells (N=171). Here juice E (O) was defined as that eliciting higher (lower) firing rate. **(d)** *Chosen value* + cells (N=56). Here trials with the two presentation orders were pooled. In the top panels, trials were sorted according to the value of offer1 (V(off1)); in the bottom panels, trials were sorted according to the value of offer2 (V(off2)). **(e)** *Chosen value* – cells (N=78). Same as in panel (e). **(f)** Untuned cells (N=757). Here we pooled cells that did not pass the ANOVA (N=729) and cells that did but were not accounted for by the selected sequences (N=28). All trials were pooled. In the top (bottom) panels, trials were sorted according to the value of offer1 (offer2). Note that the y scale is different from that used for other cell groups.

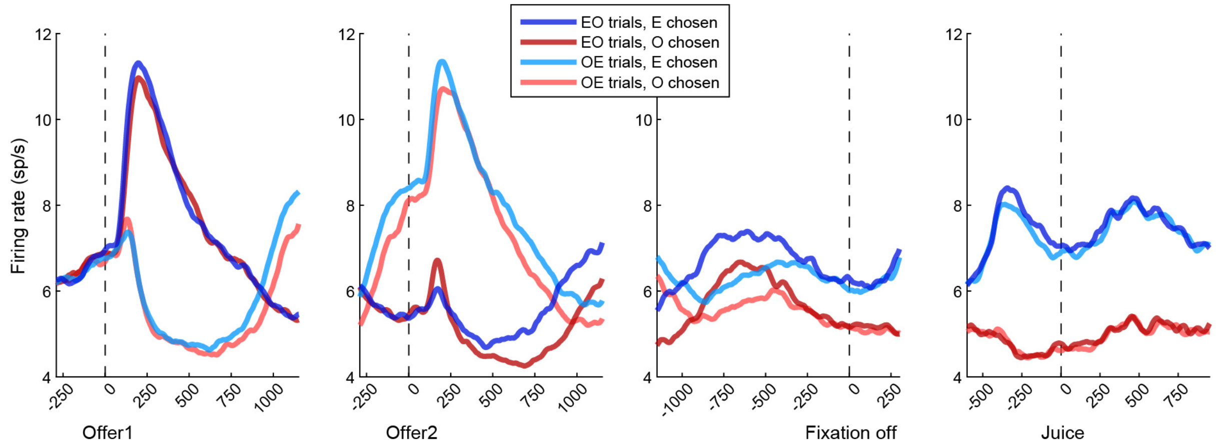


**Figure S5.** Chosen juice cells. For each neuron, we divided trials according to the presentation order (EO, OE) and the chosen juice (E chosen, O chosen). Traces shown here are averages across the population (N = 171 cells). For both EO trials and OE trials, the choice outcome signal (blue versus red traces) emerges following offer2. Interestingly, the signal is initially more pronounced for EO trials, suggesting a complex system dynamics (see Discussion).

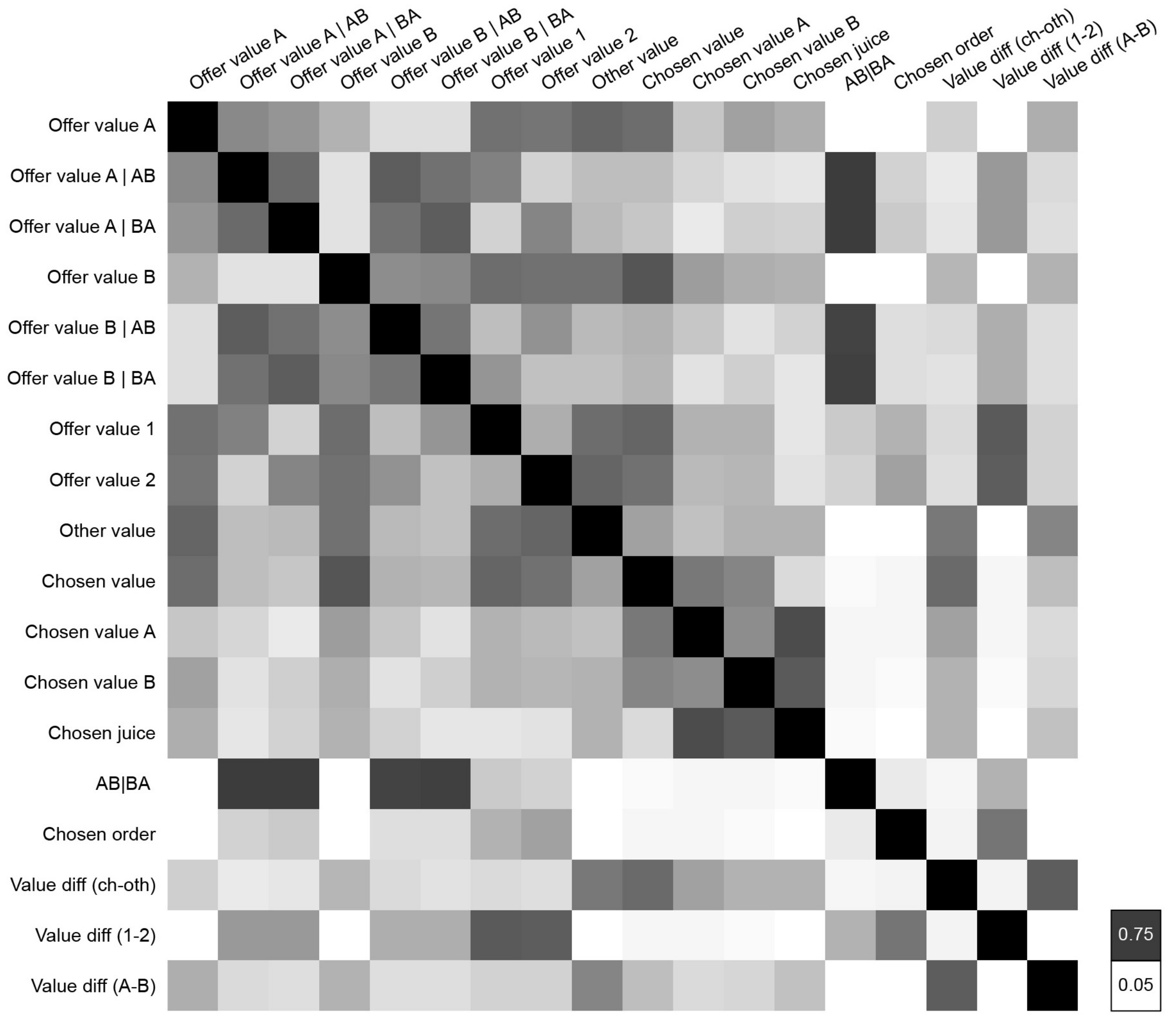


**Figure S6.** Correlation between different variables included in the analysis. The correlation matrix is rendered in gray scale such that corr = 1 is represented in black (diagonal elements) and corr = 0 is represented in white. Out of the diagonal, the maximal correlation was equal to 0.75.

| Time window | # cells |
|---|---|
| pre-offer | 4 |
| post-offer1 | 361 |
| inter-offer | 244 |
| post-offer2 | 429 |
| offers off | 270 |
| fixation off | 87 |
| pre-juice | 167 |
| post-juice | 193 |
| at least one | 612 |

**Table S1.** Results of ANOVA. A total of 1267 cells were included in the analysis. For each time window, the table indicates the number of cells that passed the ANOVA criterion ($p<0.001$). The last row indicates the number of neurons that passed the criterion in at least one of the 8 time windows.

| | Variable | Definition |
|---|---|---|
| 1 | *offer value A* | Value of juice A offered |
| 2 | *offer value A \| AB* | = *offer value A* in AB trials, 0 in BA trials |
| 3 | *offer value A \| BA* | = *offer value A* in BA trials, 0 in AB trials |
| 4 | *offer value B* | Value of juice B offered |
| 5 | *offer value B \| AB* | = *offer value A* in AB trials, 0 in BA trials |
| 6 | *offer value B \| BA* | = *offer value A* in BA trials, 0 in AB trials |
| 7 | *offer value 1* | Value of offer1 |
| 8 | *offer value 2* | Value of offer2 |
| 9 | *other value* | Value of the offer not chosen |
| 10 | *chosen value* | Value of the chosen offer |
| 11 | *chosen value A* | = *chosen value* if A chosen, 0 otherwise |
| 12 | *chosen value B* | = *chosen value* if B chosen, 0 otherwise |
| 13 | *chosen juice* | Binary; 1 if A chosen; 0 if B chosen |
| 14 | *AB \| BA* | Binary; 1 in AB trials; 0 in BA trials |
| 15 | *chosen order* | Binary; 1 if offer1 chosen; 0 if offer2 chosen |
| 16 | *value diff (ch – oth)* | = *chosen value – other value* |
| 17 | *value diff (1–2)* | = *offer value 1 – offer value 2* |
| 18 | *value diff (A–B)* | = *offer value A – offer value B* |

**Table S2.** Defined variables. Value variables were defined in units of juice B, assuming linear value functions, and based on the relative value $\rho$ derived from the logistic fit. Thus the value of $q_B$ drops of juice B was equal to $q_B$; the value of $q_A$ drops of juice A was equal to $\rho\ q_A$.

| Sequence number | post-offer1 | post-offer2 | post-juice |
|---|---|---|---|
| #1 | *offer value A \| AB +* | *offer value A \| BA +* | *chosen value A +* |
| #2 | *offer value A \| AB –* | *offer value A \| BA –* | *chosen value A –* |
| #3 | *offer value B \| BA +* | *offer value B \| AB +* | *chosen value B +* |
| #4 | *offer value B \| BA –* | *offer value B \| AB –* | *chosen value B –* |
| #5 | *AB \| BA +* | *AB \| BA –* | *chosen juice +* |
| #6 | *AB \| BA –* | *AB \| BA +* | *chosen juice –* |
| #7 | *offer value1 +* | *offer value2 +* | *chosen value +* |
| #8 | *offer value1 –* | *offer value2 –* | *chosen value –* |
| #9 | *offer value A \| AB +* | *offer value A \| BA +* | *chosen juice +* |
| #10 | *offer value A \| AB –* | *offer value A \| BA –* | *chosen juice –* |
| #11 | *offer value B \| BA +* | *offer value B \| AB +* | *chosen juice –* |
| #12 | *offer value B \| BA –* | *offer value B \| AB –* | *chosen juice +* |
| #13 | *offer value B \| BA +* | *offer value B \| AB +* | *chosen value +* |
| #14 | *offer value B \| BA +* | *offer value B \| AB +* | *chosen value –* |
| #15 | *offer value B \| BA –* | *offer value B \| AB –* | *chosen value +* |
| #16 | *offer value B \| BA –* | *offer value B \| AB –* | *chosen value –* |
| #17 | *offer value B \| BA +* | *AB \| BA +* | *chosen value +* |
| #18 | *offer value B \| BA –* | *AB \| BA –* | *chosen juice +* |
| #19 | *offer value1 +* | *offer value2 +* | *chosen juice –* |
| #20 | *offer value1 –* | *offer value2 –* | *chosen juice +* |
| #21 | *offer value1 +* | *offer value2 +* | *offer value1 +* |
| #22 | *offer value1 –* | *offer value2 –* | *offer value1 –* |
| #23 | *AB \| BA +* | *AB \| BA –* | *chosen value B –* |
| #24 | *AB \| BA –* | *AB \| BA +* | *chosen value B +* |
| #25 | *AB \| BA +* | *offer value B \| AB –* | *chosen juice +* |
| #26 | *AB \| BA –* | *offer value B \| AB +* | *chosen juice –* |

**Table S3.** Sequences that best explained at least 3 cells. Note that only few of the variables examined in the study (Table S2) ever appear in one of these 26 sequences. Sequence numbers (1st column) are arbitrary but fixed throughout the study. Collectively, these 26 sequences explained 510/538 (95%) of task-related responses.
